# Structural Insights into the Role of the Proline Rich Region in Tau Function

**DOI:** 10.1101/2024.09.20.614010

**Authors:** Karen Acosta, Christopher R. Brue, Hee Jong Kim, Polina Holubovska, Leland Mayne, Kenji Murakami, Elizabeth Rhoades

## Abstract

Tau is a microtubule-associated protein that plays an important role in modulating axonal microtubules in neurons. Intracellular tau aggregates are found in a broad class of disorders, including Alzheimer’s disease, termed tauopathies. Tau is an intrinsically disordered protein, and its structural disorder appears to be critical to its microtubule-related functions. Tubulin binding sites are found in tau’s proline-rich region (PRR), microtubule binding repeats (MTBR: R1–R4), and pseudo-repeat, R′. While many post-translational modifications have been identified on tau, phosphorylation sites, which both regulate tubulin dimer and microtubule interactions and are correlated with disease, cluster with high frequency within the PRR. Here, we use fluorescence correlation spectroscopy and structural mass spectrometry techniques to characterize the impact of phosphomimic mutations in the PRR on tubulin dimer binding and probe the structure of the PRR-tubulin dimer complex. We find that phosphomimics cumulatively diminish tubulin dimer binding and slow microtubule polymerization. Additionally, we map two ∼15 residue regions of the PRR as primary tubulin dimer binding sites and propose a model in which PRR enhances lateral interactions between tubulin dimers, complementing the longitudinal interactions observed for MTBR. Together these measurements provide insight into the previously overlooked relevance of tau’s PRR in functional interactions with tubulin.

**GRAPHICAL ABSTRACT:** 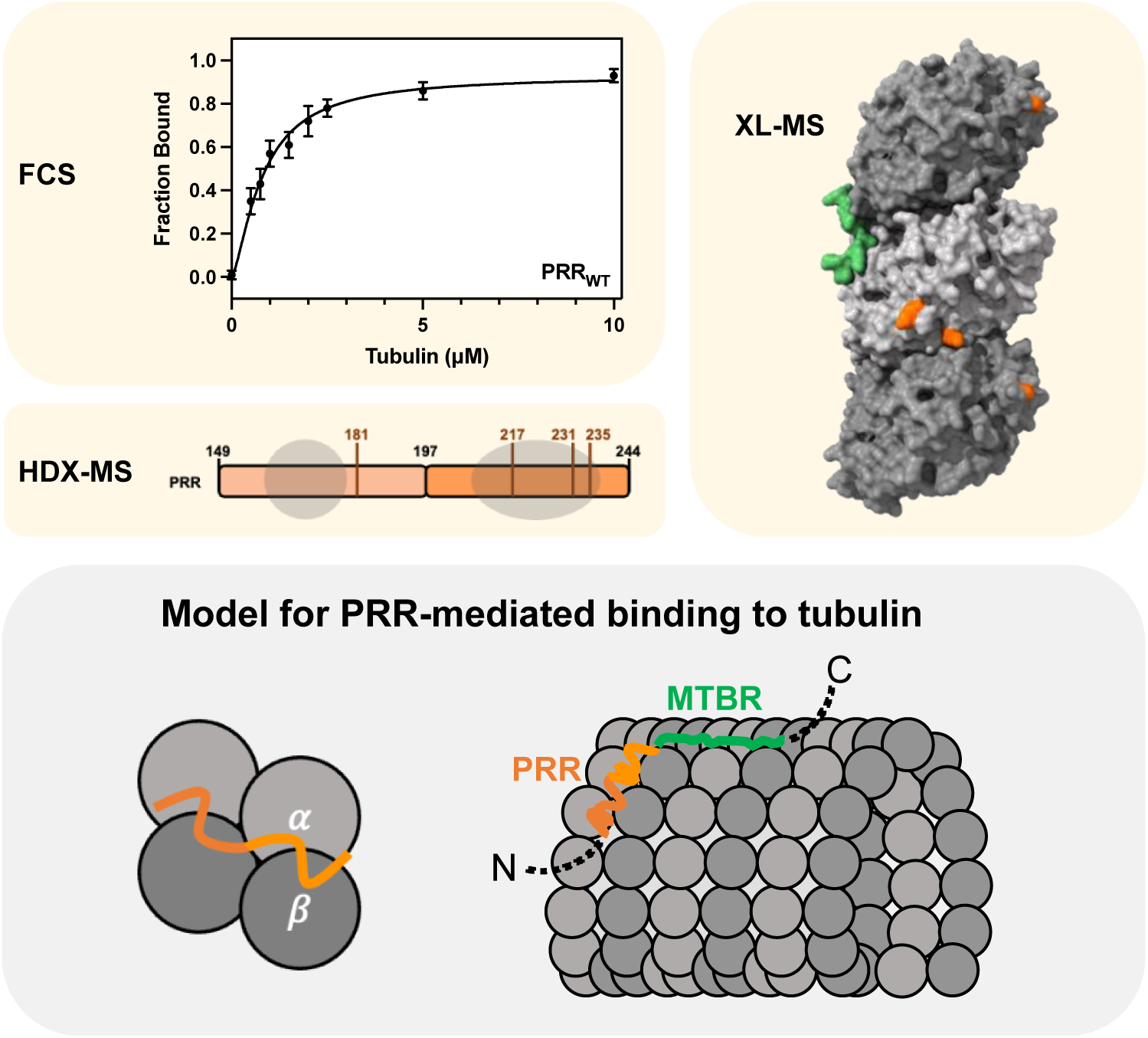

## INTRODUCTION

Tau is a microtubule-associated protein that is generally thought to function in regulating microtubule stability and dynamics^1,2^. The aggregation and deposition of tau as neurofibrillary tangles characteristic of a broad class of diseases, including Alzheimer’s disease^3^, have focused considerable attention on understanding the molecular and cellular factors that influence its self-assembly. In disease, both loss of functional tau and subsequent destabilization of microtubules, as well as gain of toxic function through aggregation, are thought to play roles in cell death^4–6^.

Tau is an intrinsically disordered protein, lacking stable secondary and tertiary structure in solution^7–9^. Even upon binding to soluble tubulin dimers ^10,11^ or microtubules^12,13^, tau remains largely disordered. Tau consists of four major domains, the N-terminal projection domain, the proline rich region (PRR), the microtubule binding region (MTBR), and the C-terminal region (Fig 1A). In the adult brain, tau is expressed as six different isoforms resulting from alternative splicing of two N-terminal inserts and R2 within the MTBR (Fig 1A). These isoforms of tau are differentially expressed based on brain regions, tissues, cell lines, intracellular compartments, and developmental stages^14–19^. Tau function is also regulated through post-translational modifications (PTM). Nearly 100 sites for PTMs have been identified on tau, including phosphorylation, acetylation, and ubiquitination, amongst others^20^. While a detailed understanding of how these numerous possible modifications impact tau function as well as its aggregation is lacking, tau phosphorylation has been heavily studied and hyperphosphorylation of tau aggregates found in post-mortem brain tissue has long been observed in disease^21^. Additionally, the phosphorylation status of tau is developmentally regulated, underscoring the importance of phosphorylation in normal tau function throughout development^22,23^. Phosphorylation patterns and frequencies may be different in the adult brain, commensurate with different requirements for microtubule growth and stability. Phosphorylation sites linked both to function and disease are found predominantly in the PRR.

**Figure 1.**
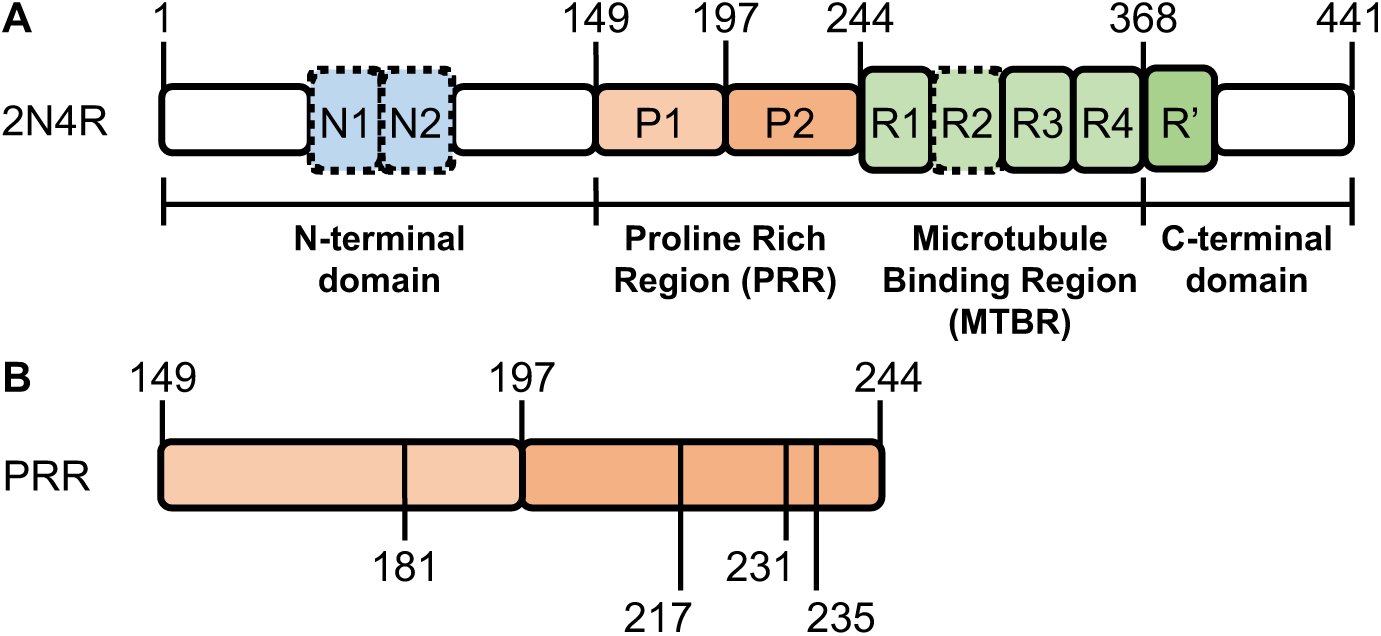
Schematic of tau with phosphomimic sites. **(A)** The longest tau isoform, 2N4R, with each of the major domains labeled: the N-terminal domain containing alternatively spliced N-terminal inserts (N1 and N2, *blue*), proline-rich region (P1 and P2, *orange*), microtubule-binding region (MTBR, *green*) with four repeat regions (R1-R4; R2 may be absent as a result of splicing), and pseudo-repeat (R′), and the C-terminal domain. **(B)** The insert highlights the PRR as well as the location of phosphomimic sites, with numbering of the residues based on 2N4R. Construct abbreviations are as follows: PRR_WT_; PRR_1P_ = PRR_T181E_ or PRR_T217E_ or PRR_T231E_ or PRR_S235E_; PRR_2P_ = PRR_T181E/T231E_ or PRR_T181E/S235E_ or PRR_T217E/T231E_ or PRR_T231E/S235E_; PRR_4P_ = PRR_T181E/T217E/T231E/S235E_.

Although PRR, MTBR, and pseudo-repeat, R′ have all been shown to be involved in interactions with microtubules^13,24–26^, the MTBR has been the focus of most intensive study, not only because of its role in binding to microtubules but also because it forms the core of aggregates in disease ^27–29^. However, recent work from our lab demonstrated that the isolated PRR was capable both of binding to soluble tubulin dimers, and had the capacity to polymerize tubulin, independent of the MTBR^30^. Intriguingly, the PRR has not been observed in structure-based models of microtubule-associated tau^12,13,31–37^. In this study, we use a combination of biophysical methods, including fluorescence correlation spectroscopy, hydrogen-deuterium exchange mass spectrometry and cross-linking mass spectrometry to provide insight into the structural features of tubulin-bound PRR. Moreover, we incorporate phosphomimic mutations (from serine or threonine to glutamate) at several locations within the PRR, both individually and in combination, to probe how modification alters this interaction. We identify regions within the PRR and sites on tubulin involved in binding and propose a model in which the PRR enhances lateral interactions between tubulin dimers.

## RESULTS

### Phosphomimics cumulatively reduce PRR binding to tubulin and microtubules

In Alzheimer’s disease patient samples, the most frequently phosphorylated sites within the PRR are T181, T217, T231 and S235^20^. As these four sites are also abundant in healthy control tissue^20^, we chose to make phosphomimic mutations from serine or threonine to glutamate both individually and in combination at each of these sites (Fig 1B). In addition, a cysteine mutation was introduced at the N-terminus (residue 149) to allow for site-specific labeling with Alexa Fluor 488 maleimide. Binding of PRR to tubulin dimers was quantified with fluorescence correlation spectroscopy (FCS) as a function of tubulin concentration. Binding to tubulin results in a shift in the PRR autocorrelation curve to longer timescales due to slower diffusion (Fig 2A)^11,30,38^. FCS measurements were made using ∼20 nM Alexa 488 labeled PRR and tubulin in concentrations ranging from 0 to 10 µM. Curves were fit to extract the fraction of PRR bound at each concentration of tubulin dimer, as described in the *Materials & Methods*. As noted previously^30^, wild-type PRR (PRR_WT_) reaches its maximum diffusion time at roughly 5 µM tubulin (Fig 2B, *black*), with no increase in binding observed even at 30 µM tubulin (Fig S1). Based on our prior published work, we know that the diffusion time at saturation corresponds to PRR bound to two tubulin dimers (Fig 2B) ^38^. For analysis of the phosphomimic variants, we fixed the diffusion time of tubulin-bound PRR to that observed for PRR_WT_ to illustrate differences between the variants (Fig 2B-F). However, as described in the *Materials & Methods*, we also evaluated other models in fitting the data; our analysis and interpretation are not dependent upon use of this model (Fig S2).

**Figure 2.**
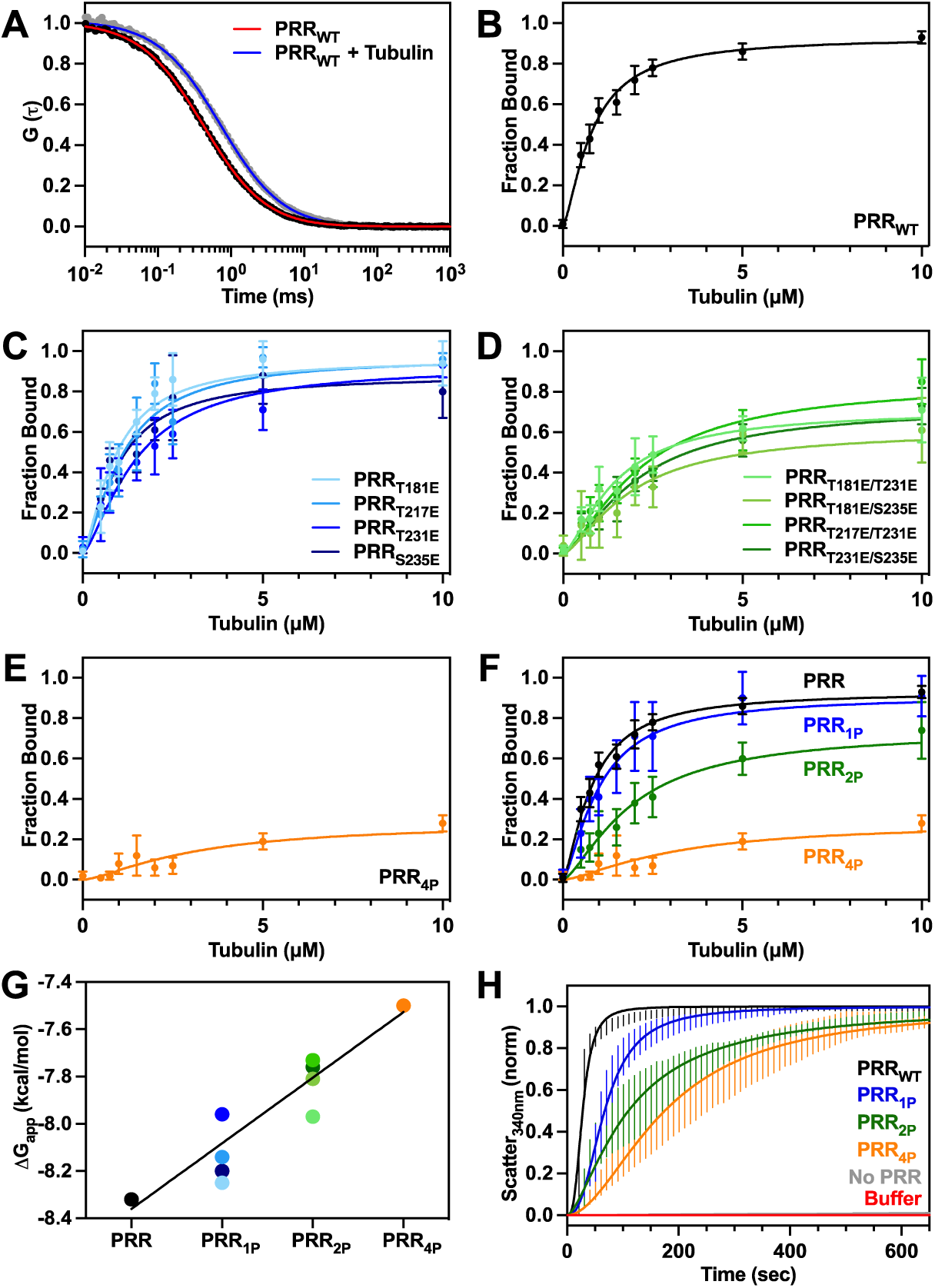
Increased phosphorylation diminishes PRR-tubulin activity. **(A)** Autocorrelation curves of free PRR_WT_ (black) and PRR_WT_ bound to tubulin dimers (gray) with two (Eqn 2) component diffusion fits for free PRR_WT_ (red) and PRR_WT_ bound to tubulin dimers (blue), respectively, as described in the *Materials & Methods*. **(B-F)** Binding of PRR constructs to tubulin dimers is quantified via fitting the fraction of tau bound as a function of tubulin concentration by the Hill equation. Data is presented as mean ± SD, for n ≥ 3 independent measurements. Constructs are represented as follows: **(B)** PRR_WT_ (black), **(C)** PRR_1P_ plotted individually (blue) **(D)** PRR_2P_ plotted individually (green) and **(E)** PRR_4P_ (orange). **(F)** PRR_1P_ (blue) and PRR_2P_ (green) were averaged to facilitate comparison amongst variants. **(G)** Relative free energy of binding, ΔG_app_ (Eqn 4), as a function of number of phosphomimics using K_D,app_ as determined by fitting to the Hill equation. Free energy was fit using a simple linear regression model (R-squared = 0.99). **(H)** PRR-induced tubulin polymerization is measured by the scattering of light at 340 nm as a function of time for 10 µM tubulin with or without 10 µM PRR construct. Data is presented as mean ± SD following normalization, for n ≥ 3 independent measurements. The polymerization data is fit with a sigmoidal equation as described in the *Materials & Methods* (solid line).

With the introduction of any individual phosphomimic, PRR_1P_, (Fig 2C, *blue*), PRR shows some reduction in tubulin dimer binding (Fig 2F). The binding curves cluster together fairly closely, with the exception of PRR_T231E_, which shows a greater reduction in binding, particularly at low tubulin concentrations, than the other single phosphomimics. It is likely that tau, and the PRR specifically, is phosphorylated at more than one site both in healthy brains as well as in the context of disease. Therefore, it is of interest to examine these effects on tubulin dimer binding of phosphomimics occurring simultaneously in the same construct. When PRR contains two phosphomimic mutations, PRR_2P_, we again observe that the binding curves cluster together (Fig 2D) and show a greater reduction in tubulin dimer binding capacity compared to PRR_WT_ and PRR_1P_ variants (Fig 2F). Although in isolation, PRR_T231E_ showed a greater diminishment in tubulin dimer binding than the other single phosphomimics, this behavior is superseded by the addition of a secondary mutation. Lastly, we measured binding by PRR with four phosphomimics present, PRR_4P_, and found it had a significantly diminished ability to bind to tubulin dimers (Fig 2E; comparison of PRR phosphomimics Fig 2F). These results suggest that, at least for the sites investigated here, that the accumulation of multiple phosphorylation events in a single construct intensify the loss of tubulin dimer binding, without a single site dominating this loss. The binding curves were fit with the Hill equation to obtain apparent dissociation constant, K_D,app_, values for comparison (Table 1). Global fitting of these binding curves yields a Hill coefficient of n = 1.5 ± 0.1^30^, in agreement with our previous measurements,^30^ emphasizing the association of PRR to more than one tubulin dimer at saturation. To estimate the relative impact of each phosphomimic on reducing the binding affinity, we converted K_D_,_app_ to an apparent free energy, ΔG_app_ (Fig 2G). We observe a linear relationship between the number of phosphomimic sites and ΔG_app_, with the addition of each phosphomimic reducing binding by ∼0.28 kcal/mol.

**Table 1.**
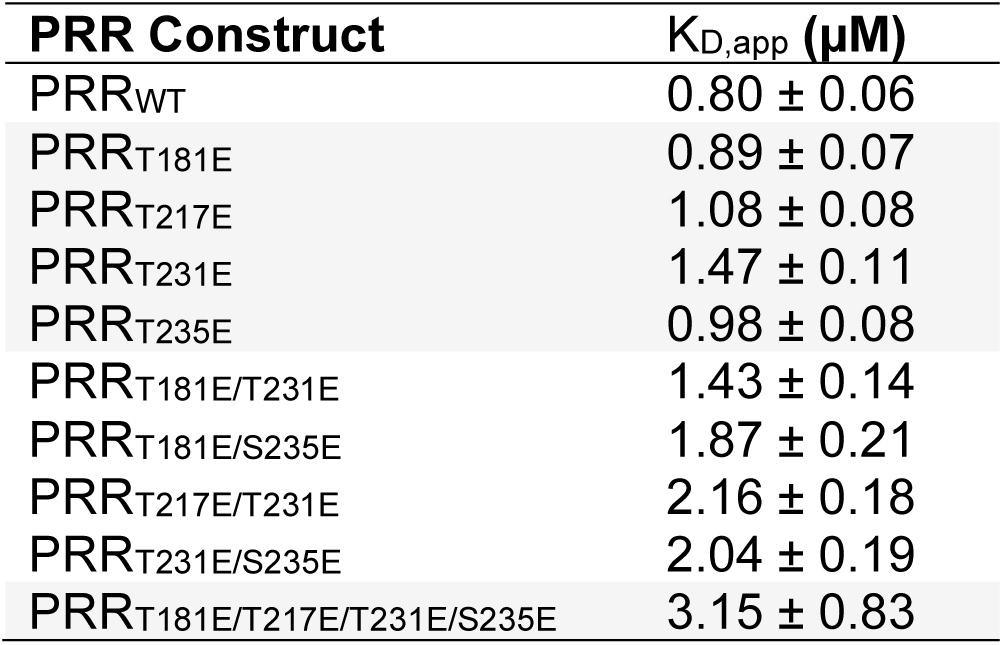
Fitting coefficients from binding curves. Binding curves derived from FCS measurements of PRR-tubulin were fit to the Hill equation, Eqn 3. The Hill coefficient was fit as a global parameter (1.5 ± 0.1), while the dissociation constants (K_D,app_), were determined individually.

| PRR Construct | $K_{D,app}$ (μM) |
| --- | --- |
| PRR<sub>WT</sub> | $0.80 \pm 0.06$ |
| PRR<sub>T181E</sub> | $0.89 \pm 0.07$ |
| PRR<sub>T217E</sub> | $1.08 \pm 0.08$ |
| PRR<sub>T231E</sub> | $1.47 \pm 0.11$ |
| PRR<sub>T235E</sub> | $0.98 \pm 0.08$ |
| PRR<sub>T181E/T231E</sub> | $1.43 \pm 0.14$ |
| PRR<sub>T181E/S235E</sub> | $1.87 \pm 0.21$ |
| PRR<sub>T217E/T231E</sub> | $2.16 \pm 0.18$ |
| PRR<sub>T231E/S235E</sub> | $2.04 \pm 0.19$ |
| PRR<sub>T181E/T217E/T231E/S235E</sub> | $3.15 \pm 0.83$ |

To investigate the impact of phosphomimics on the polymerization capacity of PRR, tubulin and PRR were co-incubated in the presence of GTP at 37°C. Polymerization was observed by an increase in light scattered and served as a proxy to track the kinetics of microtubule formation. As has been previously observed^30^, tubulin in the absence of PRR does not appreciably polymerize over the ∼10 minute timescale of these measurements (Fig 2H, *gray*). Upon the addition of PRR_WT_, the light scatter signal increases almost immediately (Fig 2H, *black*); in our previous work, we used TEM to confirm that this increase in scatter signal corresponds to microtubule formation^30^. This data is fit with a sigmoidal equation to quantify polymerization half-times (Table 2). We then compared the polymerization kinetics of PRR_WT_ with representative constructs containing one, two or four phosphomimic mutations. For PRR_1P_ the polymerization time is approximately doubled (Fig 2H, *blue*) relative to PRR_WT_. Similarly, increases in the polymerization half-times are observed for PRR_2P_ (Fig 2H, *green*) and PRR_4P_ (Fig 2H, *orange*) of approximately 4x and 6x, respectively, relative to PRR_WT_. Thus, there is correlation between the reduction in tubulin dimer binding by FCS and inhibition of microtubule polymerization capacity observed as a function of increasing numbers of phosphomimic sites. Our prior work has established that simultaneous binding of at least two tubulin dimers are required for tau fragment-mediated tubulin polymerization^38^, and this polymerization data supports this 1:2 PRR:tubulin dimer stoichiometry for PRR_4P_:tubulin complexes. While the binding of PRR_4P_ is significantly diminished relative to PRR_WT_, its tubulin polymerization capacity is retained, although the kinetics are slower. We note that this is in contrast to our previously published work comparing tubulin dimer binding and polymerization by PRR and MTBR-R’^30^; for these fragments, MTBR-R’ binding to tubulin dimers is significantly reduced compared to PRR, similar to what we observe for PRR_4P_ relative to PRR_WT_ here; however, unlike PRR_4P_, MTBR-R’ shows almost no tubulin polymerization capacity under comparable conditions.

**Table 2.** Sigmoidal fitting of microtubule polymerization assays. PRR microtubule polymerization curves were fit to Eqn 5 to determine polymerization half-times (T_50_).

| PRR Construct | T<sub>50</sub> (sec) |
| --- | --- |
| PRR<sub>WT</sub> | 27.9 ± 0.2 |
| PRR<sub>1P</sub> | 67.4 ± 0.6 |
| PRR<sub>2P</sub> | 110.0 ± 1.6 |
| PRR<sub>4P</sub> | 175.0 ± 1.4 |

While we have previously shown that the PRR is sufficient for binding to and polymerizing tubulin^30^, we had not investigated its interaction with microtubules. Both PRR_WT_ and PRR_4P_ bind to taxol-stabilized microtubules (Fig 3A), although the binding of PRR_4P_ is significantly reduced as compared to PRR_WT_ (Fig 3B). This diminishment is not due to differences in the microtubules present in each sample, as the average PRR_WT_ and PRR_4P_ intensity signal is independent of microtubule length, as is the average intensity of the microtubules (Fig S3). We note that while PRR_WT_ binds relatively uniformly to the MT lattice, PRR_4P_ binding is more variable (Fig 3A). This has also been observed for full length tau as well as PRR-containing tau fragments on microtubules and mouse hippocampal neurons ^39^. Binding of the PRR to microtubules is of particular interest, as structural models of full-length tau resolved only the MTBR bound to the microtubule lattice^12^; when combined with molecular mechanics in a metainference method, part of the PRR, P2, can also be modelled^40^. We hypothesize that the dynamic nature of PRR’s interaction with tubulin does not give rise to the stable, regular structure required for helical reconstruction in cryo-EM models. Both our PRR-tubulin dimer and microtubule measurements are also carried out at much lower PRR:tubulin ratios than cryo-EM^12^ which may allow for different binding modes.

**Figure 3.**
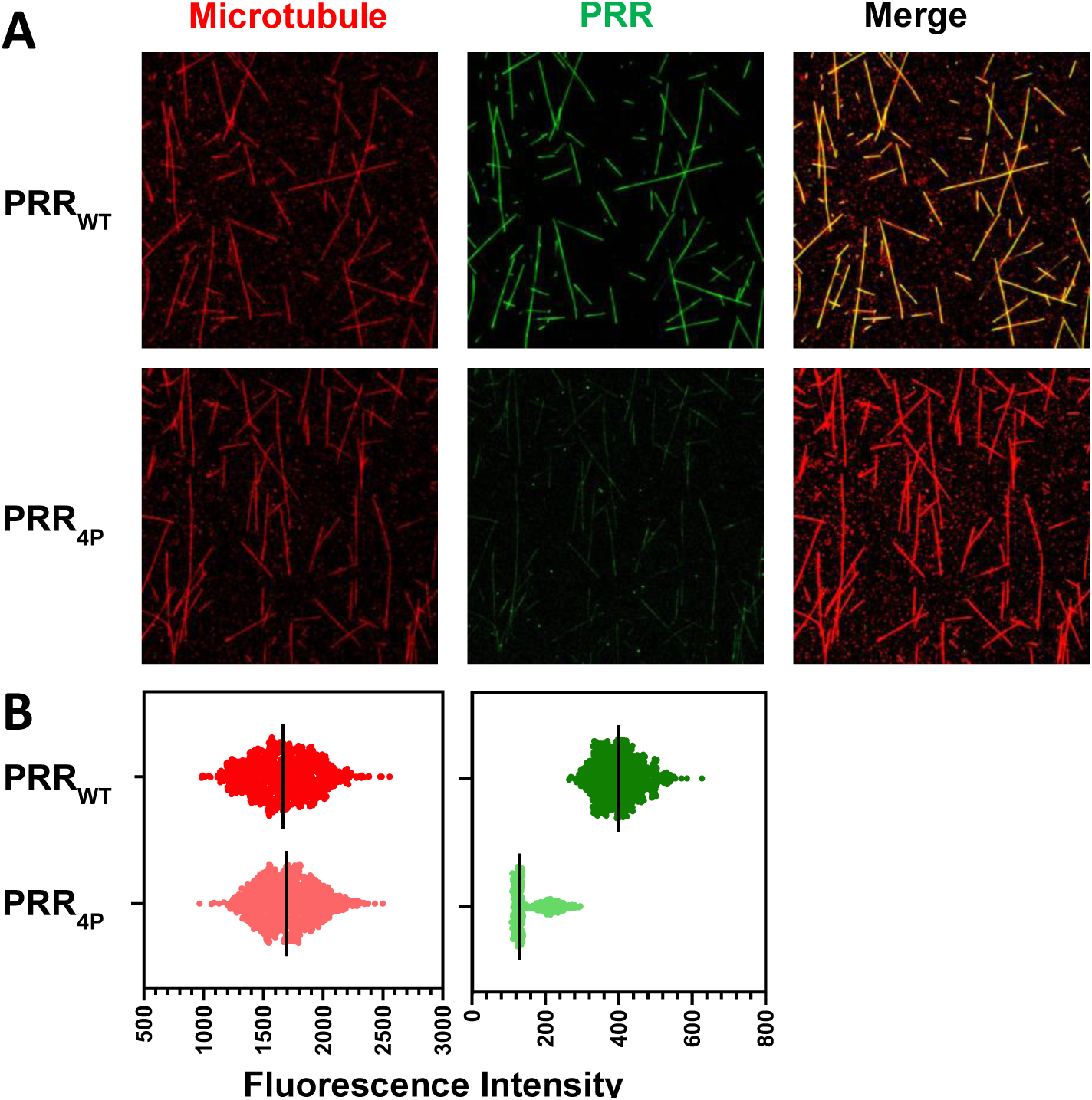
PRR binds to microtubules and phosphomimics reduce binding. **(A)** Images of microtubules (labeled with Rhodamine Red-X) co-incubated with PRR_WT_ or PRR_4P_ (labeled with Alexa 488). Labeled PRR was added to taxol-stabilized microtubules in a 0.5:1 tau:tubulin dimer ratio. **(B)** Plots of fluorescence intensity per microtubule. Each point represents the average intensity of tubulin-Rhodamine Red-X (left) or PRR-Alexa 488 (right) for a single microtubule, with the median intensity indicated by a line. N = 782 and N = 751 for PRR_WT_ microtubules and PRR_4P_ microtubules, respectively. Data represents n ≥ 3 independent measurements.

### Phosphomimics reduce dynamics for PRR alone and in the presence of tubulin

Hydrogen-deuterium exchange mass spectrometry (HDX-MS) was used to provide insight into the regions of PRR involved in tubulin binding. HDX-MS is a technique capable of resolving conformational dynamics that occur during protein binding^41^ based on changes in mass of amide hydrogens on the protein backbone upon exchange with deuteriums in solution. The rate of exchange between hydrogen and deuterium is dependent on solvent accessibility and protein structure. By coupling a commonly used pepsin column with an *Aspergillus niger* prolyl endopeptidase (AnPEP) column for PRR digest prior to mass spectrometry analysis, we were able to obtain complete coverage of the PRR, with multiple peptides for all regions (Fig S4). Our results returned digestion patterns similar to a prior application of AnPEP in HDX-MS^42^, in that we observed short peptides (>6 residues) alongside longer peptides (up to 49 residues). Shorter peptides may more accurately reflect changes in exchange than longer peptides covering the same region ^42^. As has been observed in a prior HDX-MS study of tau, deuterium exchange occurred rapidly ^43^, reaching completion for most peptides by the 30 sec timepoint (Fig S5). This timepoint was thus chosen for a point of comparison between different samples. Due to the significant diminishment in tubulin dimer binding and polymerization by PRR_4P_ relative to PRR_WT_, we chose these two constructs for comparison by HDX-MS. We identified 122 and 84 unique peptides for PRR_WT_ and PRR_4P_, respectively, with 40 of these peptides present in both samples (Fig S4). A smaller number of total peptides was identified in PRR_4P_ than in PRR_WT_ (164 as compared to 266), plausibly due to the presence of phosphomimics altering protease digestion patterns (Fig S4) or increased loss of sample due to protein aggregation (Fig S5).

HDX-MS measurements were carried out for both PRR constructs in the presence of tubulin. For PRR_WT_, tubulin dimer binding generally decreased deuterium uptake across the entire length of the protein (Fig 4A). This suggests that the much of this domain interacts with tubulin, at least transiently. Peptides covering two regions ∼165-179 and ∼224-239 reflect more enhanced protection, followed by ∼207-223; interestingly, the former is directly adjacent to the T181 phosphorylation site while the latter region encompasses the T231 and S235 phosphorylation sites. In contrast, PRR_4P_ shows much less protection in the presence of tubulin (Fig 4B). This is likely because much of the protein is not bound to the tubulin dimer, or bound only very transiently, due to its reduced affinity (Fig 2). However, the regions that show the most protection from exchange in PRR_WT_ are also amongst the more highly protected in PRR_4P_. This suggests that while the phosphomimics reduce binding of PRR to tubulin dimer, the regions involved in binding are not significantly altered.

**Figure 4.**
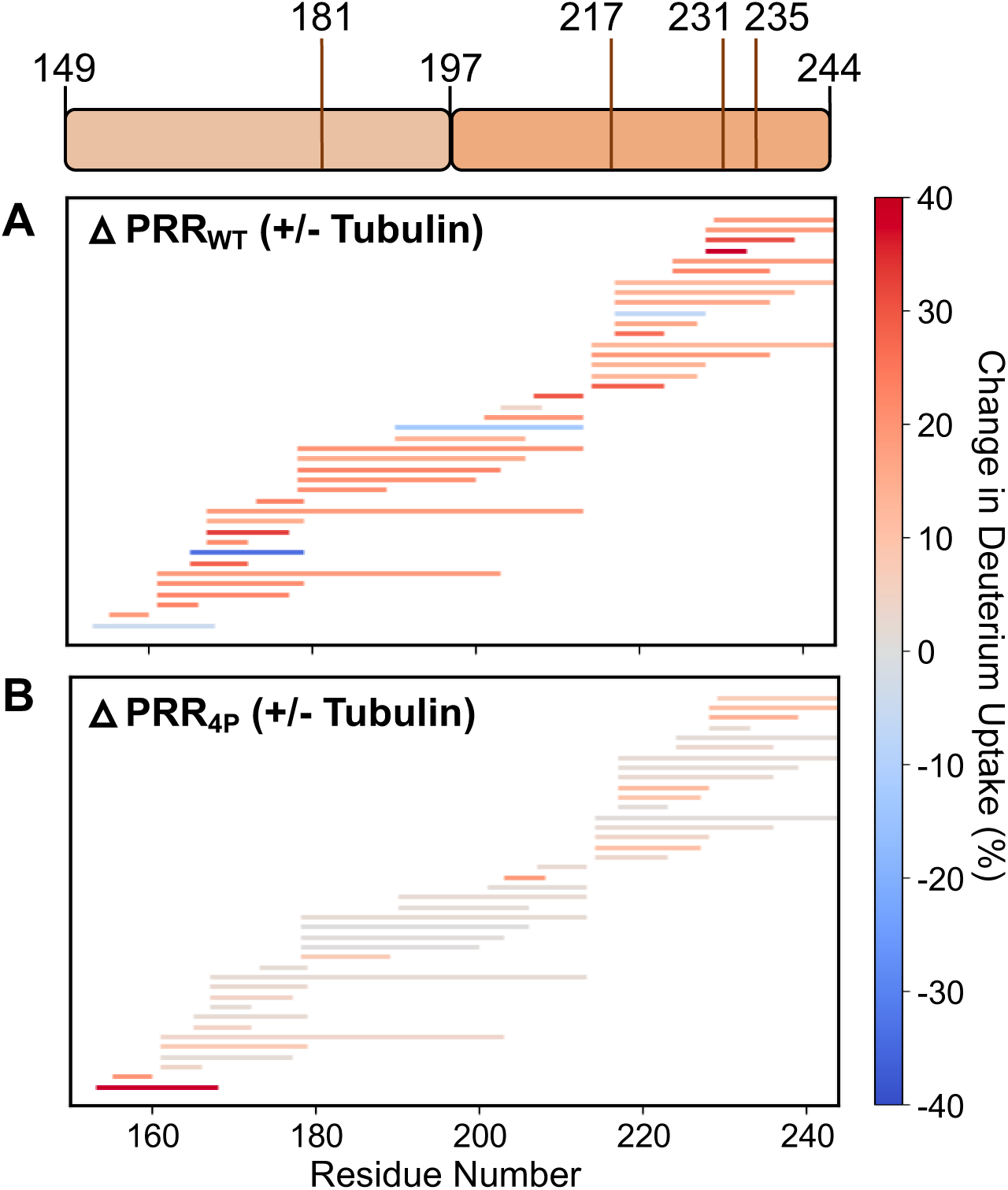
Tubulin dimer binding selectively decreases deuterium exchange in PRR. Schematic of PRR with phosphomimic sites indicated. **(A)** Map of digested PRR_WT_ or **(B)** PRR_4P_ peptides observed in HDX-MS experiments. Each horizontal bar represents a unique peptide allowed to exchange with deuterium for 30 sec identified in both PRR_WT_ and PRR_4P_ samples. The ratio of measured over maximum expected deuteriums were determined for each peptide. The horizontal bars are color coded based on the average difference in deuterium uptake reported as percent uptake. The regions shaded in gray highlight PRR residues 165-179 and 224-239. Data for each peptide is representative of n ≥ 3 independent measurements.

### PRR interacts predominantly with the **α**-tubulin subunit

Our HDX-MS measurements suggest that like the MTBR, much of the PRR interacts with the tubulin dimer surface upon binding, although dynamically. To provide further insight into tau-tubulin dimer interactions, we used cross-linking mass spectrometry (XL-MS) with the cleavable cross-linker, disuccinimidyl dibutyric urea (DSBU) to map sites of interaction between PRR and tubulin. DSBU reacts preferentially with primary amines found on lysine residues but have also been found to react with hydroxyl groups in serine, threonine and tyrosine^44^. The addition of DSBU to PRR-tubulin dimer mixtures results in a number of bands that run larger than the ∼55 kDa α/β-tubulin monomer band by SDS-PAGE (Fig S6), corresponding to various stoichiometries of ⍺/β-tubulin and PRR. We excised gel bands that were larger than the monomer tubulin band for further analysis. Cross-linked peptides were identified by mass spectrometry and identifications with a false discovery rate (FDR) of 1% or lower were not interpreted.

We collected a total of 595 cross-links containing either our PRR construct or tubulin, with 188 and 407 cross-links associated with the PRR_WT_ and PRR_4P_ reactions, respectively. The greater abundance of cross-links in the PRR_4P_ sample may in part reflect a greater number of transient interactions from the weakly-bound protein. For both samples, cross-links were observed within and between the α/β tubulin monomer units, as well as between PRR peptides (Fig 5). We primarily focus our analysis on PRR-tubulin cross-links, where there is significant overlap in the cross-linking sites found in PRR_WT_ and PRR_4P_, with K163 the most frequent cross-link position for both in the PRR, followed by K174. These residues are found directly adjacent to or within the ∼165-179 region identified by HDX-MS as protected from exchange in the presence of tubulin dimers (Fig 4). Additional sites present in both PRR_WT_ and PRR_4P_ are K224, K225 and K240. Although not as abundant as the K163 and K174 cross-links K224, K225 and K240 cross-links flank either side of the ∼224-239 region identified by HDX-MS as another putative tubulin binding site (Fig 4), and were also identified as cross-link sites between full-length tau and microtubules^33^. Interestingly, however, cross-links are notably absent at lysine residues near two of the phosphorylation sites we tested, T231/S235 (K234). This suggests that the reduced binding observed for phosphomimics at these positions may be primarily due to electrostatic repulsion^45^ rather than disruption of specific interactions with these residues and the tubulin surface. The tau:tubulin interaction has been previously shown to be electrostatically driven^13,46,47^ and FCS measurements with increasing amounts of salt demonstrate that screening charges in this way can diminish binding (Fig S7).

**Figure 5.**
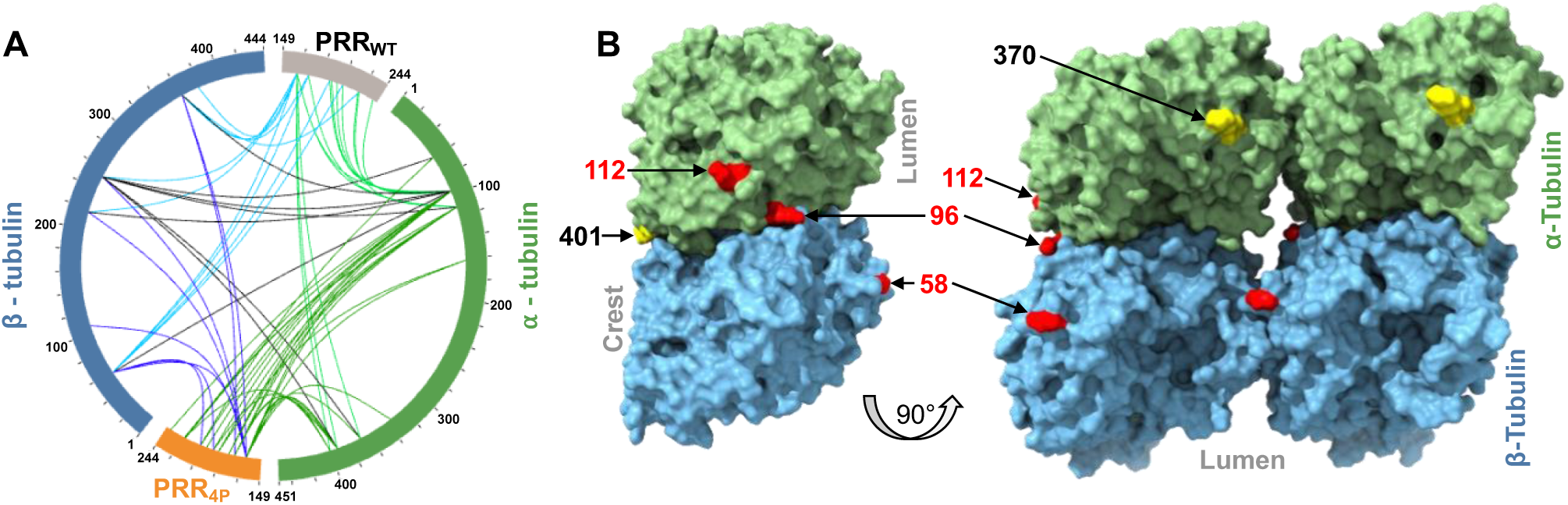
Cross-linking reveals specific PRR interactions with tubulin subunits. **(A)** Cross-links observed are shown a circle diagram visualized by xiView^50^. Cross-link colors are as follows: PRR_WT_:α-tubulin (*light green*), PRR_4P_:α-tubulin (*dark green*), PRR_WT_:β-tubulin (*light blue*), PRR_4P_:β-tubulin (*dark blue*), and α-tubulin:β-tubulin (*black*). **(B)** PRR_WT_:α/β-tubulin and PRR_4P_:α/β-tubulin cross-links were mapped on the cryo-EM structure of taxol stabilized microtubules (PDB: 6WVL) using ChimeraX 1.5^51^. The most abundant cross-link sites are colored red, with secondary sites in yellow.

PRR shows a preference for interactions with sites located on α-tubulin, with 58% of total PRR-tubulin cross-links (Fig 5). Preferential cross-linking with α-tubulin has also been observed for full-length tau^33^.The most common cross-link sites on α-tubulin were K96 and K112; however, K370, K394 and K401 were also identified in both PRR samples. On β-tubulin, K58 was the most abundant cross-linking site, with additional sites at K216 and K252 (Fig 5). We also examined the Cα–Cα distances for cross-links between α-α-, β-β-, and α-β-tubulin subunits; 97% of these found in both PRR_WT_ and PRR_4P_ samples are within 30 Å (+/-3 Å) tolerance reported for DSBU cross-linking^48^. Because tubulin is an obligate dimer, the presence of the α-β-tubulin cross-links is expected and helps corroborate our analysis. We cannot differentiate intra- or inter-molecular cross-links for α-α- and β-β-tubulin in our analysis; these may reflect PRR-driven oligomerization of the tubulin dimers^49^.

## DISCUSSION

The PRR has long been recognized for its role in enhancing tau’s interactions with microtubules^52^. More recent work from our own lab proposed that, at least under some conditions, the PRR is tau’s primary domain for both tubulin dimer-and microtubule-binding^30^. Here we use select phosphomimic mutations to probe their impact on PRR binding to soluble tubulin dimers and microtubules. Our data suggests that the accumulation of phosphomimic modifications is the primary driver for the decrease in tubulin dimer binding affinity, and subsequent decrease in tubulin-polymerization capacity. Phosphorylation of multiple sites is a common means of titrating the interaction affinity between disordered proteins and their partners^53^. This is consistent with the ‘rheostat’ model of regulation^54^, whereby each additional phosphorylation increases or decreases the free energy of the binding interaction by approximately the same magnitude (Fig 2G). The PRR contains a high fraction of the ∼55 putative phosphorylation sites that have been identified in tau^20^, and it is feasible that multiple residues will be phosphorylated concurrently to alter the functional interactions between tau and tubulin dimers or microtubules. Prior studies have shown that phosphorylation sites within the PRR have differential effects on interactions with microtubules; phosphorylation at residues 212, 214 and 231 lower the affinity for microtubules, with other sites having only minor effects^55^. Here, we find that while modification of one individual site may have a mild impact on binding affinity relative to another site, these differences are minor in comparison to the impact of multiple phosphomimic modifications simultaneously (Fig 2F). Thus, the relative strength of the interaction between tau and tubulin can be ‘tuned’ by the addition or loss of a phosphate modification, at least for the sites addressed in the current study. One recent paper used cryo-EM data combined with molecular mechanics modelling to predict phosphorylation sites that would be most destabilizing to tau-microtubule interactions^40^; interestingly, the sites identified in this study were predominantly in the MTBR, suggesting a direction for future study.

Disordered proteins often interact with binding partners through short stretches of residues or short linear motifs or SLiMs^56^. Our HDX-MS measurements identify two stretches of residues within the PRR that are more protected from exchange in the PRR-tubulin complex, ∼165-179 and ∼224-239 (Fig 4). Interestingly, this latter region encompasses a stretch of residues first identified by Feinstein and coworkers more than 25 years ago^52^, 224-230 (identical to 215-221 in their study) as strongly influencing microtubule binding and assembly in fragments of tau containing the MTBR. A more recent NMR study of full-length tau also identified roughly the same two regions of the PRR showing increased loss of signal relative to flanking regions upon binding both the soluble tubulin dimers and microtubules^33^. For PRR_WT_, an increase in protection is seen throughout the PRR sequence in the presence of tubulin, suggesting that much of the protein associates, at least transiently, with the tubulin dimer surface (Fig 4A). This is similar to binding models also proposed for the MTBR – while specific residues are important for binding, the entire region associates along the microtubule lattice. In contrast, PRR_4P_ shows little protection from exchange outside of the primary binding sites (Fig 4B) suggesting that PRR_4P_ is not stably associated with tubulin dimers over the timescale of the HDX-MS measurements. Even though HDX-MS reports on the average behavior of the complex, this is supported by our FCS data which also indicates faster off-rates for PRR_4P_.

The XL-MS results provide further insight into PRR binding sites on tubulin dimers and, importantly, how these may differ with microtubules. In particular, some tubulin residues available in soluble tubulin are obscured by neighboring tubulin dimers in microtubules. Of the cross-links identified in this study, K370 on α-tubulin and K58 on β-tubulin each face the luminal side of a microtubule, while K96 on α-tubulin and K58 on β-tubulin are buried by lateral or longitudinal contacts between dimers (Fig 5). These tubulin sites that are not accessible in microtubules may reflect the interactions important for initial association of PRR with soluble tubulin dimers to initiate polymerization. The cross-linked sites found on the opposite sides of α-tubulin subunit, K112 and K401, are accessible on the outside of the microtubule (Fig 5). Moreover, single molecule FRET measures ∼90 Å RMS of PRR_WT_ bound to tubulin ^30^ approximately the span of two laterally adjacent tubulin subunits. Collectively, our data leads us to propose a model whereby PRR stabilizes lateral interactions between tubulin dimers (Fig 5). This is in contrast to the structural model derived from full-length tau, where the MTBR was found to bind longitudinally along the crest of the microtubule protofilament^12^. It is of note that the tubulin binding sites identified for PRR in this study do not overlap with the binding sites of MTBR such that both regions could be bound simultaneously (Fig 5). This also allows for the possibility of conditions where MTBR anchors tau to the microtubule, while PRR plays a role in attracting tubulin dimers or other binding partners to the microtubule lattice.

In summary, this study provides insight into the impact of phosphomimic mutations on interactions between PRR and tubulin. We find that phosphomimics have a cumulative effect on reducing binding to tubulin dimers and microtubules. The PRR-tubulin complex is dynamic, driven primarily by interactions between sequences embedded within the PRR and electrostatic attraction between negative tubulin and positive PRR; and the introduction of phosphomimics or phosphorylation, likely diminishes these favorable interactions. Lastly, we propose a model whereby PRR enhances lateral interactions between tubulin dimers, with minimal overlap between its interaction with tubulin and the MTBR binding site (Fig 6). It would be of interest to investigate if the nucleotide state of tubulin/microtubules alters PRR binding patterns ^57^ as well as what role the disordered tubulin tails play in PRR-mediated binding and polymerization. Our study emphasizes the power of a combination of biophysical approaches to provide critical insights into the molecular details and structural features of a dynamic complex which may be challenging to characterize by traditional structural approaches.

**Figure 6.**
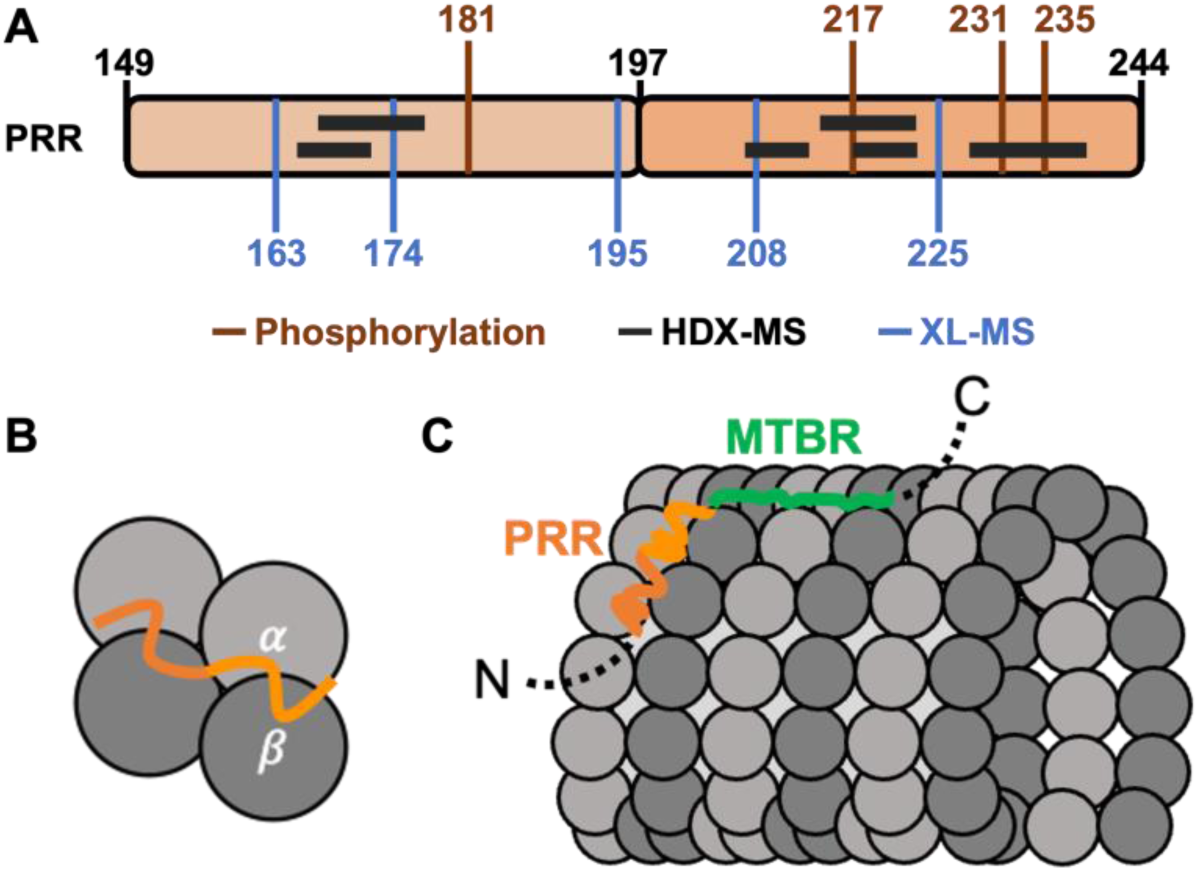
Model for PRR-tubulin interactions in full-length tau. **(A)** PRR schematic split into two segments for P1 and P2, with phosphorylation sites indicated. Regions of highest protection from exchange and top cross-link sites are indicated. Model of tau with PRR (orange) bound to **(B)** two tubulin dimers or **(C)** microtubules with MTBR (green) along the crest of the protofilament. α-tubulin and β-tubulin are represented in light gray and dark gray respectively.

## ACKNOWLEDGMENTS

We thank JASCO Facility for Spectroscopic Excellence for support of the fluorimeter and the Biological Chemistry Resource Center for use of the ultracentrifuges, both in the Department of Chemistry at the University of Pennsylvania. Hydrogen-deuterium exchange mass spectrometry measurements were performed at the Johnson Foundation Biophysics and Structural Biology Core at the University of Pennsylvania. Cross-linking experiments were supported in part by the NIH, grant number R01GM123233 (to KM). This work is funded by the NIH, grant number RF1AG053951 (to ER), the Structural Biology and Molecular Biophysics Training Grant NIH T32 GM008275 (to KA and PH) and Training in Age Related Neurodegenerative Diseases T32AG000255 (to CB).

## MATERIALS AND METHODS

### Tubulin purification

Tubulin was purified as previously described^58^ from young bovine brains with two cycles of polymerization and depolymerization with one modification: the concentration of nucleotides was adjusted to 1.5 mM ATP and 1 mM GTP for the first polymerization, then increased to 2.5 mM ATP and 1.5 mM GTP for the second polymerization step. The resulting pure tubulin was flash frozen in liquid nitrogen in BRB80 (80 mM PIPES, 1 mM MgCl_2_, 1 mM EGTA, pH 6.8) and stored at -80 °C. As needed, aliquots were thawed quickly at room temperature, then clarified via centrifugation at 100,000xg for 6 min at 4 °C. BioSpin 6 columns (BioRad) were used to exchange tubulin into the desired buffer. The tubulin concentration was determined by using the absorbance at 280 nm and molar extinction coefficient of 115,000 M^-1^cm^-1^, and used within 2 hours of clarification.

### Fluorescent labeling of tubulin

The tubulin labeling protocol was adapted from a previously published protocol^59^. To 200 µL tubulin (5-10 mg/mL), 22 µL 5X BRB80, 1.6 µL 500 mM MgCl_2_ and 2.24 µL 100 mM GTP were added, and the solution was incubated on ice for 5 minutes. To this solution, 100 µL 37 °C glycerol was added and mixed well by pipetting. The solution was incubated in a 37 °C water bath for 30 minutes to polymerize tubulin. The tubulin mixture was layered onto 200 µL 37 °C high pH cushion (0.1 M HEPES, pH 8.6, 1 mM MgCl_2_,1mM EGTA, 60% v/v glycerol). The microtubules were pelleted by centrifugation at 192,000xg for 45 minutes at 35 °C. The supernatant was removed by pipette aspiration and discarded, followed by gentle rinsing of the supernatant-cushion interface twice with labeling buffer (0.1 M HEPES, pH 8.6, 1 mM MgCl_2_,1mM EGTA, 40% v/v glycerol). All liquid, including the cushion, was removed by pipette aspiration. The tube was returned to the 37 °C water bath and 120 µL labeling buffer, also at 37 °C, was added. The pellet was resuspended by pipetting, using a pipette tip with the end cut off to avoid shearing the microtubules.

Assuming ∼70% recovery of initial tubulin, Rhodamine Red-X succinimidyl ester (from a 13 mM stock in anhydrous DMSO) was added to 10-fold molar excess to the 120 µL microtubule solution. The microtubule-dye solution was incubated for 30-40 minutes in a 37 °C water bath, with vortexing for 5 seconds every 2 minutes. 120 µL quench buffer (2X BRB80, 100 mM K-glutamate, 40% v/v glycerol) was added and mixed well by pipetting. The solution was incubated in a 37 °C water bath for 15 minutes, the pipetted onto 180 µL low pH cushion (1X BRB80 pH 6.9, 60% v/v glycerol). The solution was centrifuged at 300,000xg for 20 minutes at 37 °C.

The supernatant was removed by pipette aspiration and discarded. The supernatant-cushion interface was washed twice with warm BRB80 and discarded. All remaining liquid, including the cushion, was removed by pipetting and 60 µL ice cold BRB80 was added to the pellet. The pellet was resuspended by pipetting, using a pipette tip with the end cut off. Pipetting was continued until no chunks were visible. The solution was incubated on ice for at least 60 minutes, then centrifuged in a pre-chilled rotor at 300,000xg for 10 minutes at 2 °C. The supernatant was recovered, and the final tubulin concentration and labeling stoichiometry were determined by absorbance.

### PRR expression and purification

The proline rich domain (PRR, tau amino acids 149-244) was cloned into a pET-HT vector containing an N-terminal HisX6 tag with a tobacco etch virus (TEV) cleavage site. The plasmid was transformed into BL21(DE3) cells and grown in 0.5 L LB media with ampicillin (100 μg/mL) until reaching absorbance at 600 nm of 0.5-0.6. At this point, expression was induced by the addition of 1 mM IPTG for 4-6 hours at 37 °C. The cells were pelleted and resuspended in 50 mM Tris pH 8.0, 500 mM NaCl, 10 mM imidazole, 1 mM PMSF, and one cOmplete protease inhibitor cocktail tablet. The lysed cells were flash frozen in liquid nitrogen and stored at -80°C until use. To thawed lysate, 1 mg/mL of lysozyme was added prior to sonication. The sonicated lysate was then centrifuged at 20,000xg for 20 minutes at 4 °C to remove cellular debris. The supernatant was syringe filtered through a 0.22 μm filter and incubated with ∼5-7 mL of Ni-NTA resin for 1 hour at 4 °C with rotation. The beads were washed with Buffer A (50 mM Tris pH 8.0, 500 mM NaCl, 10 mM imidazole), and PRR eluted in a single step with Buffer B (50 mM Tris pH 8.0, 500 mM NaCl, 400 mM imidazole). The eluate was concentrated and buffer exchanged into Buffer A using Amicon Ultra Ultracel-3K. The protein was then incubated with TEV protease at 4 °C overnight with rocking. The cleaved protein was once again incubated for 1 hour in Ni-NTA beads at 4 °C with rotation. The cleaved PRR was collected in the flow-through, while the cleaved N-terminal HisX6 tag and any remaining uncleaved PRR and the TEV protease all remained associated with the Ni-NTA beads. The PRR was concentrated and buffer exchanged into Buffer C (25 mM Tris pH 8, 100 mM NaCl, 1 mM EDTA, 1 mM TCEP) using Amicon Ultra Ultracel-3K. The sample was filtered as before and loaded on Superdex 200 HiLoad 16/600 for further purification. Purified protein was flash frozen in liquid nitrogen and stored at -80 °C until use.

### Fluorescent labeling of PRR

For site-specific labeling of PRR, a cysteine (PRR contains no native cysteines) was introduced at the N-terminus, residue 149. Expression and purification of this construct was as described above. For labeling, ∼300 μM PRR (determined by UV-Vis with ε_280_ = 1,490 M^-1^cm^-1^) was treated with 1 mM DTT and incubated at room temperature for 30 mins to reduce the cysteine. Amicon concentrators were used as described previously to remove the DTT and exchange into labeling buffer (20 mM Tris pH 7.4, 50 mM NaCl) with 6M Guanidine HCl. For labeling, Alexa Fluor 488 maleimide (AL488) was added in 3-5X molar excess to protein and incubated overnight with stirring at 4 °C. Prior to its use in the labeling reaction, AL488 was dissolved in anyhdrous DMSO, aliquoted and flash frozen in liquid nitrogen for storage at -80 °C. For labeling, a single aliquot was quickly thawed prior to a labeling reaction, with any unused dye discarded. Any remaining unconjugated dye and Guanidine HCl were removed by buffer exchange into labeling buffer with Amicon concentrators, followed by passing the sample through two coupled 5 mL HiTrap desalting columns equilibrated with labeling buffer. Labeled proteins were aliquoted, flash frozen in liquid nitrogen, and stored at -80 °C until use.

### FCS instrumentation and analysis

All FCS measurements were performed on our home-built instrument as described previously ^10^. The laser (488 nm diode-pumped solid-state laser, Spectra-Physics) was adjusted to ∼5 μW prior to entering the inverted Olympus 1X-71 Microscope (Olympus) and focused using a 60x 1.2 NA water immersion objective. Fluorescence emission was collected through the objective and separated from excitation light by a Z488RDC long pass dichroic and a 500LP long pass filter (Chroma). Collected emission was focused onto the aperture of a 50 μm diameter optical fiber (OzOptics) directly coupled to an avalanche photodiode (Perkin-Elmer). A digital correlator (FLEX03LQ-12, Correlator.com) was used to calculate the autocorrelation curves. Alignment of instrument and analysis were verified using Alexa Fluor 488 hydrazine (Fisher).

Measurements were made in 8-chamber coverslips (Nunc, Lab-Tek) passivated by incubation with polylysine conjugated to polyethylene glycol (PEG-PLL) to reduce protein adsorption^11^. For each measurement, ∼15-25 nM labeled PRR was incubated with tubulin (a range of concentrations was used) for 5 min in Nunc chambers in phosphate buffer (20 mM phosphate buffer pH 7.4, 20 mM KCl, 1 mM MgCl_2_, 0.5 mM EGTA, and 1 mM DTT) at 20 °C. Each measurement consisted of 25 traces of 10 seconds. The autocorrelation curves were averaged and fit to a one-component (Eqn 1: PRR alone) or a two-component (Eqn 2: PRR + tubulin) 3D diffusion equation using lab-written scripts in MATLAB (Mathworks). These measurements were run in triplicate on different days and averaged to obtain statistical variations.

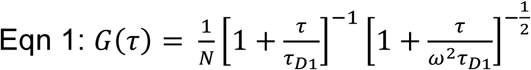

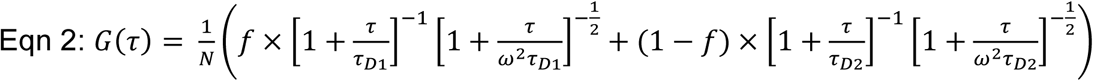

G(τ) is the autocorrelation function as a function of time, where τ_*D*1_ and τ_*D*2_ are the diffusion times of free PRR_WT_ and bound PRR_WT_ with 10 μM tubulin, respectively; N is the average number of fluorescent species in the focal volume; *f* is the fraction of free PRR; and s is the ratio of the radial to axial dimensions of the focal volume. For our instrument, s was determined to be 0.2 using Alexa 488 hydrazine (Invitrogen) as a reference standard and subsequently fixed for all fitting. We note that the apparent decrease in the fraction bound plateau seen in Fig 2 for PRR_2P_ and PRR_4P_ likely reflects dynamic exchange of tubulin in the PRR-tubulin complex as it diffuses through the observation volume, resulting in a decrease in the apparent τ_*D*2_ of those constructs^60^. We thus chose to use the τ_*D*2_ of PRR_WT_ at 10 μM tubulin for fitting of all FCS curves to underscore differences in binding by the phosphomimics for Fig 2.

Binding curves were constructed by plotting the fraction bound, f_b_=(1-f) from the FCS fits as a function of tubulin, which were then fit to the Hill equation (Eqn 3).

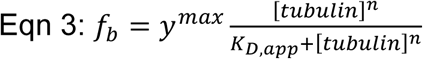

The curves were fit with the Hill coefficient (n) as a global parameter, *y^max^* fixed to the maximum value of f_b_ for each construct, and the apparent dissociation constants (K_D,app_) as individual parameters.

In order to test the dependence of the apparent dissociation constants on the model used for analyzing the FCS data, we also fit the FCS curves using two additional models: (1) the single diffusing component equation (Eqn 1), where τ_D_ reflects an average of the free and bound PRR species; (2) the two diffusing components equation (Eqn 2) but determining τ_*D*2_ at 10 μM tubulin for each phosphomimic construct independently, rather than using a single τ_*D*2_ determined for PRR_WT_. The binding curves derived from these fits are shown in Fig S2 along with the apparent dissociation constants, K_D,app_, for all three fit models (Table S1). These data show that the relative differences in apparent affinity/avidity are only weakly dependent upon the exact fitting model used for the FCS data. The apparent free energy, ΔG_app_ plotted in Fig 2G was calculated as:

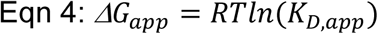

R is the gas constant, 0.001987 kcal/mol*K, T is 298.15 K, and K_D_,_app_ values from Table 1.

### Microtubule polymerization assay

Polymerization of soluble tubulin dimers into microtubules was monitored through the increase of scattered light at 340 nm as described previously ^30^. Tubulin aliquots were clarified via centrifugation and buffer exchanged into phosphate buffer (20 mM phosphate buffer pH 7.4, 20 mM KCl, 1 mM MgCl_2_, 0.5 mM EGTA, and 1 mM DTT). For each reaction 10 μM tubulin was incubated with 10 μM PRR for 2.5 minutes on ice. After the addition of 1 mM GTP, the reaction was immediately transferred to a cuvette and measured for 10 min at 37 °C in a fluorometer (FP-8300 with temperature accessory ETC-815, JASCO) with excitation and emission at 340 nm. Thereafter, the samples were returned at 4 °C for 5 min to establish the absence of protein aggregation with cold depolymerization. These measurements were run in triplicate and averaged to obtain statistical variations. Polymerization curves were fit to a sigmoidal equation (Eqn 5).

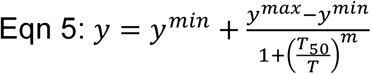

In which y^min^ and y^max^ are the minimum and maximum values for the signal of light at 340 nm over time (T) in seconds to obtain half-times (T_50_). The slope (m) is the light scattering of y^max^_-ymin._

### Preparation of taxol-stabilized microtubules

Taxol stabilized microtubules were made following a previously published protocol, with minor adaptions^61^. All tubulin solutions were thawed on ice. Tubulin (38 µM), Rhodamine Red-X tubulin (1 µM) and biotin tubulin (1 µM) were added to final volume of 20 µL in BRB80. The solution was pipetted to mix and then clarified by centrifugation at 100,000xg for 6 minutes at 4 °C.

The supernatant was removed to a new ultracentrifuge tube and 1 µL GTP (from a 20 mM stock in water) was added. The solution was incubated in a 37 °C water bath for 20 minutes to polymerize tubulin. Following incubation, 50 µM taxol in BRB80 was added in 2 µL increments for a total of 8 µL. Following each incremental taxol addition, the tube was flicked several times to mix. The solution was incubated in a 37 °C water bath for 15 minutes to polymerize the tubulin. The microtubules were pelleted by centrifugation in a pre-warmed rotor at 353,000xg for 5 minutes at 37 °C. Following centrifugation, the supernatant was removed and discarded, and the pellet was resuspended in 10 µL of 20 µM taxol in BRB80.

### Preparation of sample chambers for TIRF imaging

Coverslips and glass slides were cleaned according to a previously published protocol^62^. Briefly, coverslips and glass slides were wiped with Kimwipes then placed in a holding chamber filled with 100% acetone and incubated for 1 hour. A 10 minute incubation in 100% ethanol followed, to remove all traces of acetone, followed by two 5 minutes incubations in milli-Q water. Next, the coverslips and glass slides were incubated in freshly prepared 100 mM potassium hydroxide for 15 water, then dried with Kimwipes and stored.

The clean coverslips were placed into a coplin jar with a sufficient volume of 2% dimethyldichlorosilane to completely submerge them. After a 5 minute incubation, the coverslips were removed, and excess silane was rinsed off with two washes of 100% ethanol of 5 minutes each. Lastly, the coverslips were incubated three times, 5 minutes each, with milli-Q water. The resulting silanized coverslips were stored in a dedicated box separated by lens paper. Chambers of approximately 5 µL capacity were created just prior to use using strips of double-sided tape, silanized coverslips and cleaned glass slides.

To functionalize the surface, a 1% neutravidin solution in phosphate buffered saline was added into the chamber and incubated for 10 minutes. The chamber was rinsed with 10 volumes of phosphate buffer (20 mM phosphate buffer pH 7.4, 20 mM KCl, 1 mM MgCl2, 0.5 mM EGTA, and 1 mM DTT). Two volumes of a 5% Pluronic F-127 solution in phosphate buffer were added to the chamber and incubated for 10 minutes. The chambers were rinsed with 10 volumes of phosphate buffer and left filled with buffer until ready for use.

### Imaging of PRR-Microtubules by TIRF

Taxol stabilized microtubules (described above) were diluted 1:400 in phosphate buffer supplemented with 20 µM taxol. PRR_WT_ or PRR_4P_ labeled with Alexa 488 was added to a final 1:4 molar ratio with the microtubules (in tubulin concentration units). The microtubule/PRR solutions were incubated for 10 minutes and then loaded into sample chambers. The chambers were sealed with CoverGrip to prevent evaporation prior to imaging.

Imaging of PRR/microtubules by TIRF microscopy was conducted at ambient temperature on an inverted Eclipse Ti2 microscope (Nikon) with 100x Apo TIRF objective lens (1.49 NA) and sCMOS camera, running NIS Elements version 5.11.01. The PRR was imaged by excitation with a 488nm laser (green channel) while the microtubules were excited with a 561 nm laser (red channel). All images were obtained with a 1 s exposure. For each laser, the same illumination power was used for all images to allow for comparison.

TIRF microscopy images were analyzed using FIJI (ImageJ; National Institutes of Health). Overlapping areas of uniform illumination within the red and green channels were determined and only microtubules within this area were analyzed. Microtubules were randomly selected from the red channel images; selected microtubules were copied directly into the corresponding green channel image using the ROI Manager feature to select for microtubule bound PRR. For each microtubule, the length and average brightness was calculated in both the green and red channels. For the analysis, 782 (PRR_WT_) or 751 (PRR_4P_) microtubules from 33 total images obtained from n ≥ 3 independent sample preparations were analyzed.

### Prolyl endopeptidase purification

*Aspergillus niger* Prolyl endopeptidase (AnPEP) was purified from dietary supplement (GliadinX) by dissolving one capsule in 2 mL of Buffer D (100 mM phosphate pH 4, 100 mM NaCl). The mixture was syringe filtered through a 0.22 μm filter and 1 mL of the solution was loaded onto and eluted from a Superdex 200 HiLoad 16/600 size exclusion column. Fractions containing AnPEP were pooled, concentrated and buffer exchanged into Buffer E (100 mM phosphate pH 8, 20 mM NaCl) using Amicon Ultra Ultracel-30K. The sample was filtered as before and loaded on a 5 mL HiTrap Q HP. For separation, a 10-70% Buffer F (100 mM phosphate pH 8, 1 M NaCl) gradient was applied over 25 column volumes. AnPEP eluted at approximately 300 mM salt. This was repeated with the second 1 mL of the filtered enzyme solution before ultimately pooling relevant fractions post anion exchange. Purified AnPEP was buffer exchanged into Buffer D and stored at 4 °C overnight prior to coupling to column resin. The concentration of AnPEP was determined by using the absorbance at 280 nm, molar extinction coefficient of 125,305 M^-1^cm^-1^, with molecular weight of 72.1 kDa and final yield of ∼50 mg per capsule (Fig S8).

### Prolyl endopeptidase coupling

AnPEP was coupled to POROS-20AL (Fisher) resin by forming covalent bonds between primary protein amines and the reactive aldehyde groups on the column resin. To prepare the column, 0.2 g of POROS-20AL resin was added to 100 mg of AnPEP in 1 mL of Buffer D (100 mM phosphate pH 4, 100 mM NaCl). 320 μL of Buffer H (100 mM sodium citrate pH 4.4, 1.5 M sodium sulphate) and 100 μL of Buffer K (1 M sodium cyanoborohydride) were added slowly. Then, a total of 660 μL of Buffer H was added over a 2 hr period (27.5 μL every 5 min), with the solution placed on a rocker at room temperature between additions. The mixture was incubated overnight at room temperature with continuous rocking. The reaction was quenched with 150 μL of Buffer I (100 mM sodium citrate pH 4.4, 0.1 M ethanolamine, 5 mM sodium cyanoborohydride) and then placed back on the rocker for 1 hr at room temperature. Using a sintered glass funnel the resin was washed with 12 mL of Buffer G (100 mM sodium citrate pH 4.4), 10 mL Buffer J (100 mM sodium citrate pH 4.4, 1 M NaCl), then 12 mL of Buffer G. The resin was resuspended in a 50:50 solution (v/v) of resin:Buffer G and immediately used to prepare a column for HDX-MS. Any remaining resin was stored at 4 °C until use.

### HDX-MS peptide list preparation

For initial peptide list generation 10 µM PRR_WT_ (or PRR_4P_) in phosphate buffer (100 mM NaPO4, 150 mM NaCl, pH 7.45) was injected on to a home-built system for combined protease degradation and subsequent LC-MS^2^ identification of peptide fragments(Kan et al., 2019)1. Briefly, the injected sample passed through two columns packed with protease-loaded resin (as described above; proteases used are pepsin and AnPEP) for 4 min at 0 ^0^C. The resulting peptides were desalted on a C8 trap column. After desalting, peptides were eluted through an analytical C8 column and introduced to a Thermo Q Exactive ESI-MS. The elution profile of the analytical separation was a H_2_O/ACN gradient of 5%-40% over 15 mins. LC-MS^2^ data was analyzed with Proteome Discover (v. 2.4 Thermo-Fisher) to detect and identify peptide fragments based on the initial sequence of the PRR constructs. The initial list was then used to generate reported peptide coverage maps (Fig S4) were generated using EXMS2 analysis software ^63,64^.

### HDX-MS sample preparation data acquisition

For each exchange reaction concentrated PRR was diluted 4x in deuterated exchange buffer (100 mM NaPO4, 150 mM NaCl, pD 7.45) and allowed to exchange at RT for 10 to 6000s (Fig S5). A 30 s exchange was chosen for the analysis in this manuscript, as longer timepoints often showed significant loss of peptides. The reaction was quenched at 0 °C via 2x dilution in deuterated quench buffer (100 mM NaPO4, 150 mM NaCl, 2M Guanidine HCl, pD = 2.5l). The final concentration of PRR in all HDX reactions was 10 µM. “All-H” control samples were prepared in the same manner, but standard protonated formulations of the above exchange and deuterated buffers were used. Samples HDX were analyzed in the same manner as those used for peptide list generation except MS^1^ spectrometry was conducted to detect to the total mass difference in each detected peptide. For the back exchange corrected samples (Fig S9), samples were prepared in same manner except the samples were incubated at 50°C for 30 mins before quenching. For measurements in the presence of tubulin, PRR_WT_ (or PRR_4P_) was combined in a 2:1 (tubulin dimer:PRR) molar ratio in phosphate buffer (100 mM NaPO_4_, 150 mM NaCl, pH = 7.45) and incubated for 3 min at room temperature. The mixture was then diluted 4x in deuterated exchange buffer and analyzed in the same manner as PRR-only samples.

### HDX-MS data analysis

Deuterated peptides from each HDX-MS sample were detected and analyzed using EXMS2 software as previously published ^63^. Briefly, each HDX-MS sample was compared to the “All-H” protonated to determine the retention time of each peptide generated from the protease digest. Changes in m/z ratio of each peptide are interpreted as incorporation of deuterons. Peptide maps showing comparative deuteration were generated using python (v 3.11), pandas (2.1.2), and matplotlib (v 3.8.0). Percent deuteration was determined was determined by comparing the observed mass change to the maximum possible deuteration sites. Coloration on peptide maps is represented according to this percentage.

Since the same peptides were used for analysis in the presence and absence of tubulin (Fig 4), the relative changes in exchange should not depend upon a correction for back-exchange. Except for illustrative purposes (Fig S9), peptides were not corrected for back-exchange. The back exchange corrected deuterium uptake (in number of deuterons) for each peptide (D_corr_) was calculated as in Eqn 6: where m is the observed centroid mass at time t, m_0%_ is the centroid mass for the non-deuterated control, and m_100%_ is the centroid mass of the 100% deuterated, or maximally labeled sample. D_corr_ is represented as a percentage of the total possible amide backbone exchange sites in each peptide (Fig S9).

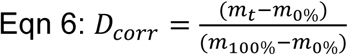

### Cross-linking mass spectrometry sample preparation

50 μg of purified tau and 50 μg of purified tubulin was mixed with 6 mM DSBU (Thermo) in phosphate buffer (20 mM phosphate buffer pH 7.4, 20 mM KCl, 1 mM MgCl2, 0.5 mM EGTA, and 1 mM DTT) and incubated on ice for 1 hour. The reaction was quenched by adding 50 mM ammonium bicarbonate (AB) then incubated on ice for 30 min. The cross-linked material was separated by SDS-PAGE (Fig S6) and gel bands were cut into ca. 1 mm^3^ pieces. The gel pieces were destained in 30% (v/v) ethanol at 60 °C for 30 min, dehydrated with 100% acetonitrile (ACN) at room temperature for 1 hour, and dried in a laminar flow hood overnight. Gel pieces were stored at −20°C for analysis by MS.

### XL-MS sample preparation

Gel pieces were rehydrated and dehydrated a few more times with 25 mM AB and ACN respectively. Gel pieces were incubated with 500 μl of 25 mM AB with final 20 ng/μl trypsin (Thermo Fisher Scientific) at room temperature overnight. Digested peptides were recovered by incubating gel pieces with 400 μl 60% ACN, 0.1% (v/v) trifluoroacetic acid (TFA; Thermo Fisher Scientific) for 10 min in the shaker (3x). The peptides were then evaporated to dryness and resuspended with 0.1% (v/v) TFA to fractionate peptides with PierceTM High pH Reversed-Phase Peptide Fractionation Kit (Thermo Fisher Scientific) according to the recommended protocol by the supplier. All fractions were vacuum-dried and resuspended with LC-MS grade water containing 0.1% (v/v) TFA for MS analysis.

Each fraction was analyzed by a Q-Exactive HF mass spectrometer (Thermo Fisher Scientific) coupled to a Dionex Ultimate 3000 UHPLC system (Thermo Fisher Scientific) equipped with an in-house–made 15-cm-long fused silica capillary column (75 μm inner diameter), packed with a reversed-phase ReproSil-Pur C18-AQ 2.4-μm resin (Dr. Maisch GmbH, Ammerbuch, Germany) column. Elution was performed using a gradient from 5 to 45% B (90 min), followed by 9% B (5 min), and re-equilibration from 90 to 5% B (5 min) with a flow rate of 3-400 nl/min (mobile phase A: water with 0.1% formic acid; mobile phase B: 8 % ACN with 0.1% formic acid). Data were acquired in data-dependent tandem MS (MS/MS) mode. Full-scan MS settings were as follows: mass range, 300 to 1800 (mass/charge ratio); resolution, 120,000; MS1 AGC target 1^E6^; MS1 Maximum IT, 200 ms. MS/MS settings were as follows: resolution, 30,000; AGC target 2^E5^; MS2 Maximum IT, 300 ms; fragmentation was enforced by higher-energy collisional dissociation with stepped collision energy of 25, 27, 30; loop count, top 12; isolation window, 1.5 m/z; fixed first mass, 130; MS2 Minimum AGC target, 800; charge exclusion: unassigned, 1, 2, 3, 8 and > 8; peptide match, off; exclude isotope, on; dynamic exclusion, 45 s^65^.

### Cross-linked peptide search

Raw files were converted to mzML format with ThermoRawFileParser 1.2.3^66^. Search engine MeroX 2.0.1.4^67^ was used to identify and validate cross-linked peptides. FASTA file was generated manually with proteins of interest entries with Uniprot *Escherichia coli* (strain B / BL21-DE3) proteome and EBI Reference proteome of *Bos taurus* (Uniprot Release 2021_03). MeroX was run in RISEUP mode, with default cross-linker mass and fragmentation parameters for DSBU with minor tweaks: precursor mass range, 1000 to 10,000 Da; minimum precursor charge, 4; precursor and fragment ion precisions, 5.0 and 10.0 ppm, respectively; maximum number of missed cleavages, 3; oxidation of methionine, as variable modification; results were filtered for score (>10) and target-decoy approach FDR (<1 %). All search results from each fraction’s MS acquisition were combined and filtered by recalculated FDR at 1 %. K-K residue pairs were selected for the site-localization when cross-linked peptides contained miscleaved lysine residues and site localization scores were tie. Visualization of the cross-links were performed with xiView^50^ and ChimeraX 1.5^51^.

