## SUPPLEMENTAL INFORMATION for "Structural Insights into the Role of the Proline Rich Region in Tau Function"

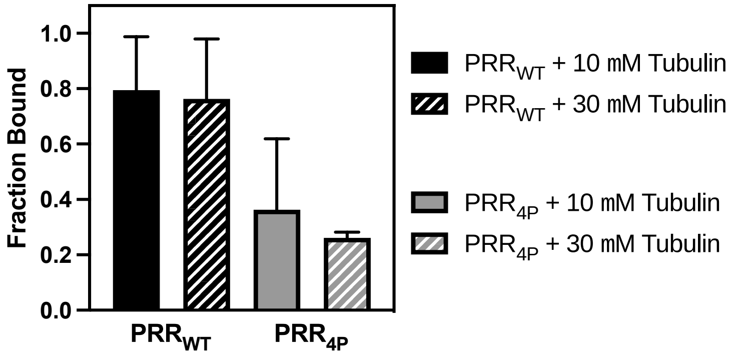


**Figure S1. 10 µM tubulin dimer is sufficient for binding measurements.** Binding of PRR constructs to tubulin dimers represented as the fraction tau bound as a function of tubulin concentration. $PRR_{WT}$ (black) and $PRR_{4P}$ (gray) with 10 µM tubulin dimer (solid) and 30 µM tubulin dimer (striped). Even with 3x more tubulin dimer, no increase in binding is calculated, indicating binding is maximal at 10 µM tubulin dimer. The 10 µM data points are used in all analysis in the main manuscript and is shown here for ease of comparison with the 30 µM tubulin data points. Data is presented as mean ± SD, for n ≥ 3 independent measurements.

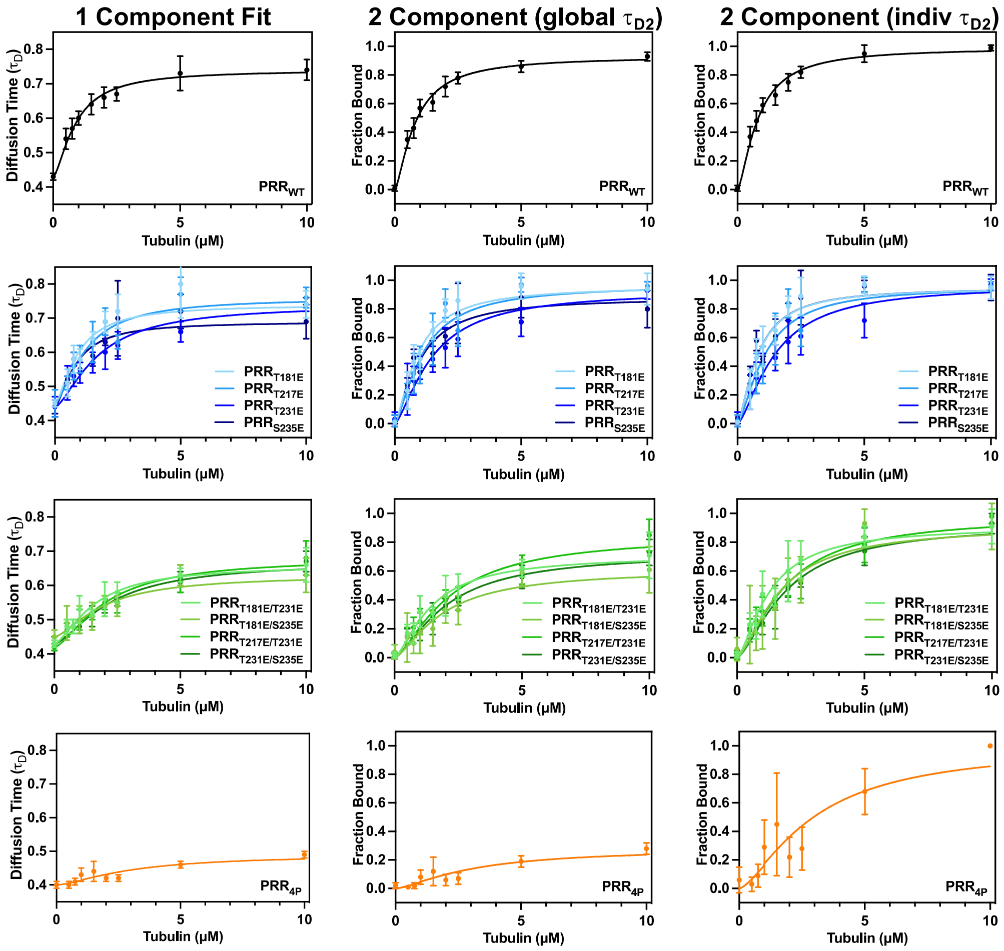


**Figure S2. Comparison of different models for fitting FCS curves.** Binding curves for PRR to tubulin dimers derived from the two alternative fitting methods described in the *Materials & Methods*: Eqn 1, where $\tau_{D1}$ reflects an average diffusion of free and bound PRR (left); Eqn 2, where the bound diffusion time, $\tau_{D2}$, was determined by the maximum diffusion time for $PRR_{WT}$ (center; reproduced from Fig 2 in *Results* to allow for comparison) or independently for each phosphomimic construct (right). Constructs are represented as follows: $PRR_{WT}$ (black), $PRR_{1P}$ plotted individually (blue), $PRR_{2P}$ plotted individually (green) and $PRR_{4P}$ (orange). The Hill equation (Eqn 3) was used to extract an apparent dissociation constant, $K_{D,app}$, from either the diffusion time (Eqn 1) or the fraction bound (Eqn 2) as described in the text. The Hill coefficient was fit as a global parameter (1.5 ± 0.1), while the dissociation constants ($K_{D,app}$ ; Table S1), were determined individually as described in the *Materials and Methods*. Data is presented as mean ± SD, for n ≥ 3 independent measurements.

| **PRR Construct** | 1 Component<br>**$K_{D,app}$ (μM)** | 2 Component (global $\tau_{D2}$)<br>**$K_{D,app}$ (μM)** | 2 Component (indiv $\tau_{D2}$)<br>**$K_{D,app}$ (μM)** |
|---|---|---|---|
| $PRR_{WT}$ | 0.88 ± 0.08 | 0.80 ± 0.06 | 0.80 ± 0.08 |
| $PRR_{T181E}$ | 0.90 ± 0.08 | 0.89 ± 0.07 | 0.79 ± 0.08 |
| $PRR_{T217E}$ | 1.20 ± 0.10 | 1.08 ± 0.08 | 1.07 ± 0.10 |
| $PRR_{T231E}$ | 1.71 ± 0.15 | 1.47 ± 0.11 | 1.48 ± 0.14 |
| $PRR_{T235E}$ | 0.82 ± 0.09 | 0.98 ± 0.08 | 0.79 ± 0.08 |
| $PRR_{T181E/T231E}$ | 1.58 ± 0.18 | 1.43 ± 0.14 | 1.27 ± 0.12 |
| $PRR_{T181E/S235E}$ | 1.93 ± 0.28 | 1.87 ± 0.21 | 1.65 ± 0.16 |
| $PRR_{T217E/T231E}$ | 2.03 ± 0.20 | 2.16 ± 0.18 | 1.86 ± 0.17 |
| $PRR_{T231E/S235E}$ | 2.23 ± 0.25 | 2.04 ± 0.19 | 1.93 ± 0.19 |
| $PRR_{T181E/T217E/T231E/S235E}$ | 3.11 ± 0.97 | 3.15 ± 0.83 | 3.01 ± 0.29 |

**Table S1. Comparison of $K_{D,app}$ from different FCS fitting approaches.** The Hill equation (Eqn 3) was used to extract an apparent dissociation constant, $K_{D,app}$, from either the diffusion time (Eqn 1) or the fraction bound (Eqn 2, global or individual $\tau_{D2}$) as depicted in Fig S2. The Hill coefficient was fit as a global parameter (1.5 ± 0.1), while the dissociation constants were determined individually as described in the *Materials and Methods*. Column 2 is from the analysis described in the *Results*; these values are found separately in Table 1 and are reproduced here to allow for comparison. Data is presented for n ≥ 3 independent measurements.

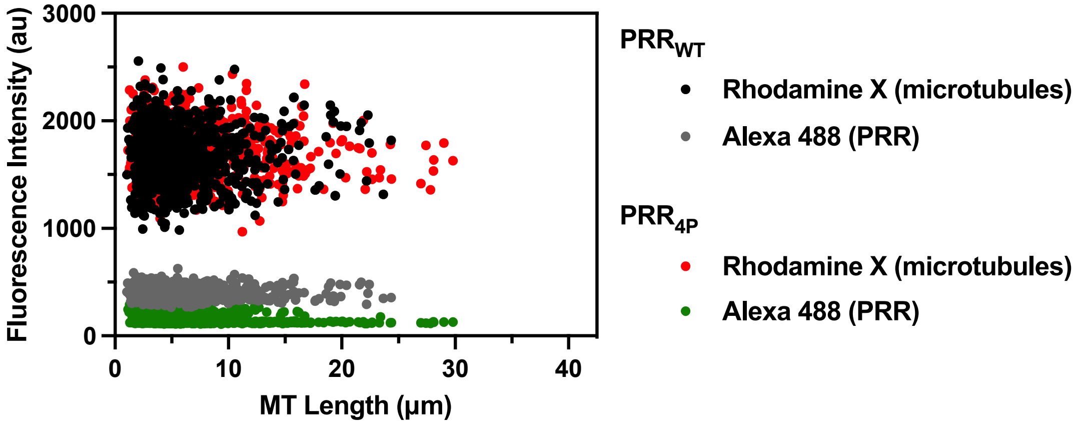


**Figure S3. Fluorescence intensities are independent of microtubule length.** Plot of average fluorescence intensities of microtubules (black, red) and PRR (gray, green) as a function of measured microtubule length for individual microtubules.

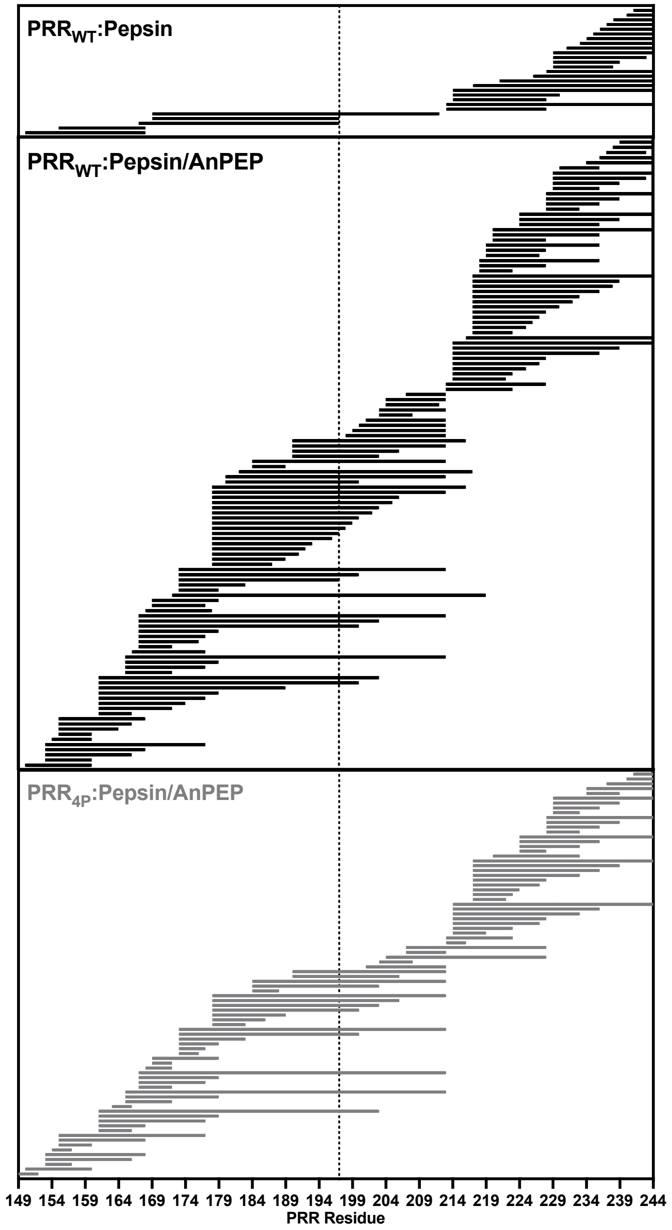


**Figure S4. Peptide coverage for $PRR_{WT}$ and $PRR_{4P}$ in HDX-MS experiments.** Each horizontal bar represents a unique peptide identified. The dotted line delineates P1 and P2 of $PRR_{WT}$ (black) or $PRR_{4P}$ (gray) as indicated. Proteins were digested with either pepsin alone or tandem pepsin/AnPEP proteases as described in the *Materials and Methods*. The inclusion of the AnPEP column significantly enhances the number of peptides obtained relative to the more commonly used pepsin column (compare the top and middle maps), resulting in full coverage of the sequence.

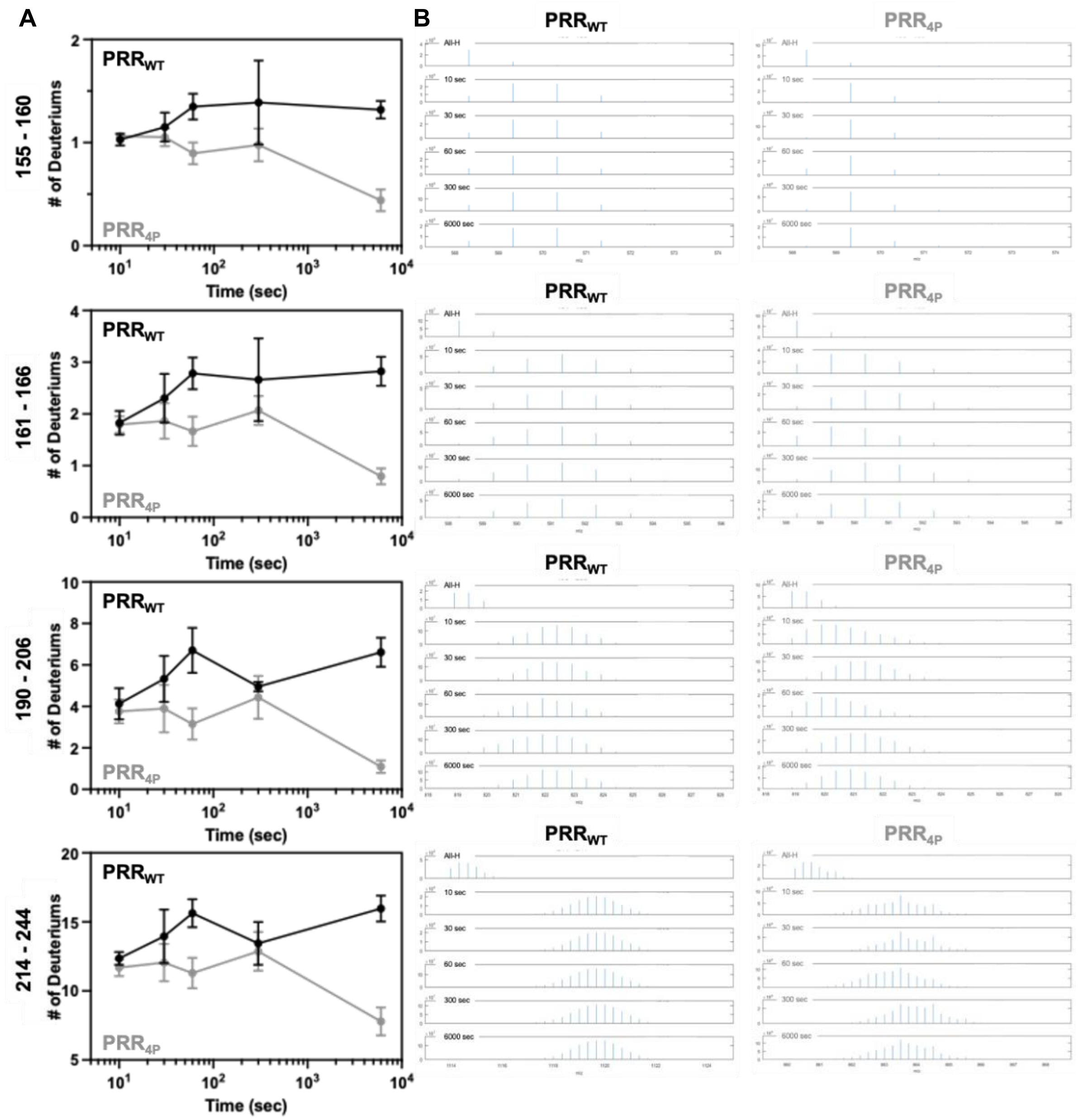


**Figure S5. Kinetic deuterium uptake profiles for selected peptides. (A)** Deuterium uptake measurements for $PRR_{WT}$ (black) or $PRR_{4P}$ (gray) at several time points. Samples showed increased variability and, in particular for $PRR_{4P}$, increased loss of peptides at longer time points, likely due to aggregation. Data is presented as mean ± SD, for n ≥ 3 independent measurements. **(B)** Corresponding mass spectra for $PRR_{WT}$ or $PRR_{4P}$ in the absence of tubulin for selected peptides at several time points, and in the absence of deuterium (All-H). The envelopes show a shift to higher masses due to deuteration following dilution into deuterium buffer.

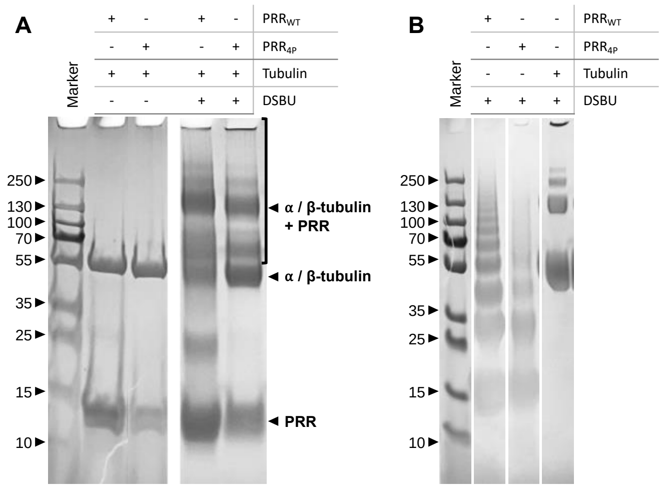


**Figure S6. SDS-PAGE of cross-linked PRR-tubulin.** Gel bands indicated by a bracket were excised and submitted to XL-MS. **(A)** Both $PRR_{WT}$ and $PRR_{4P}$ form high molecular weight assemblies with tubulin upon the addition of cross-linker DSBU as evident through SDS-PAGE. **(B)** Cross-linking controls of each protein $PRR_{WT}$, $PRR_{4P}$, or tubulin with itself.

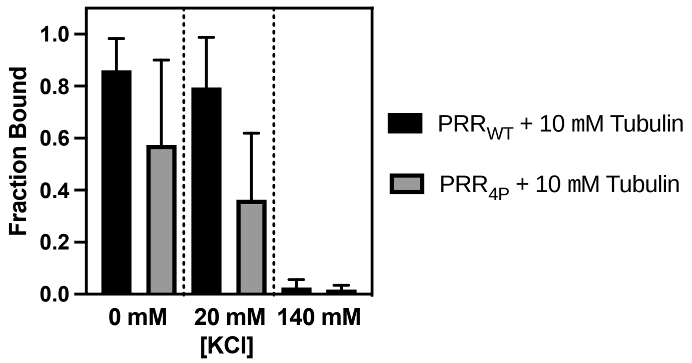


**Figure S7. Salt effects on PRR:tubulin binding.** Binding of PRR constructs to tubulin dimers represented as the fraction tau bound as a function of tubulin concentration. Constructs are represented as follows: $PRR_{WT}$ (black) and $PRR_{4P}$ (gray) with 10 µM tubulin in 0 mM, 20 mM and 140 mM salt buffer conditions. Data is presented as mean ± SD, for n ≥ 3 independent measurements.

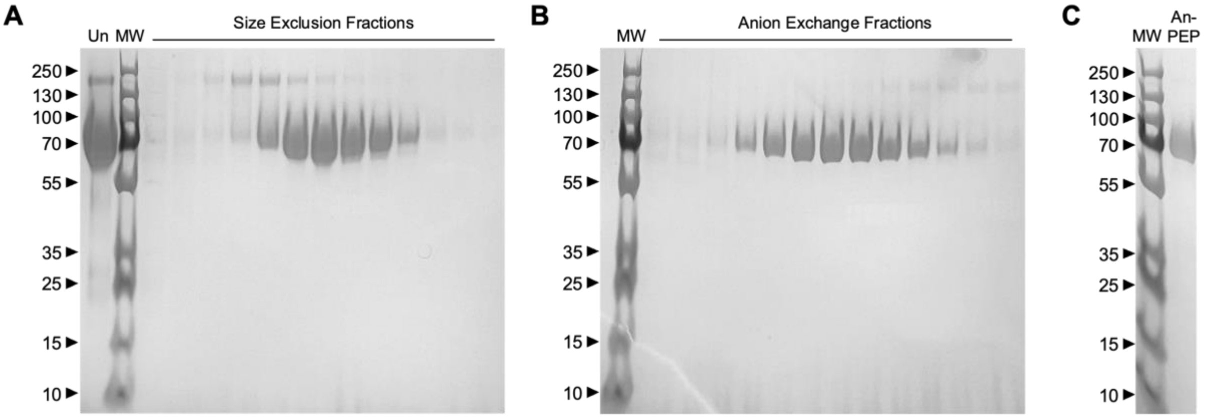


**Figure S8. Purification of Prolyl Endopeptidase.** Prolyl endopeptidase (AnPEP) was purified from dietary supplement (GliadinX, AVI Research) by dissolving the contents of one capsule in 2 mL 100 mM phosphate pH 4, 100 mM NaCl. **(A)** The initial solution (Un) was further purified by size exclusion column (Superdex 200 HiLoad 16/600). Fractions containing protein were pooled, concentrated and buffer exchanged into 100 mM phosphate pH 8, 20 mM NaCl. **(B)** Remaining impurities were separated from AnPEP by elution from an anion exchange column (HiTrap Q HP) using a 1 M NaCl gradient. **(C)** Purified AnPEP from one capsule.

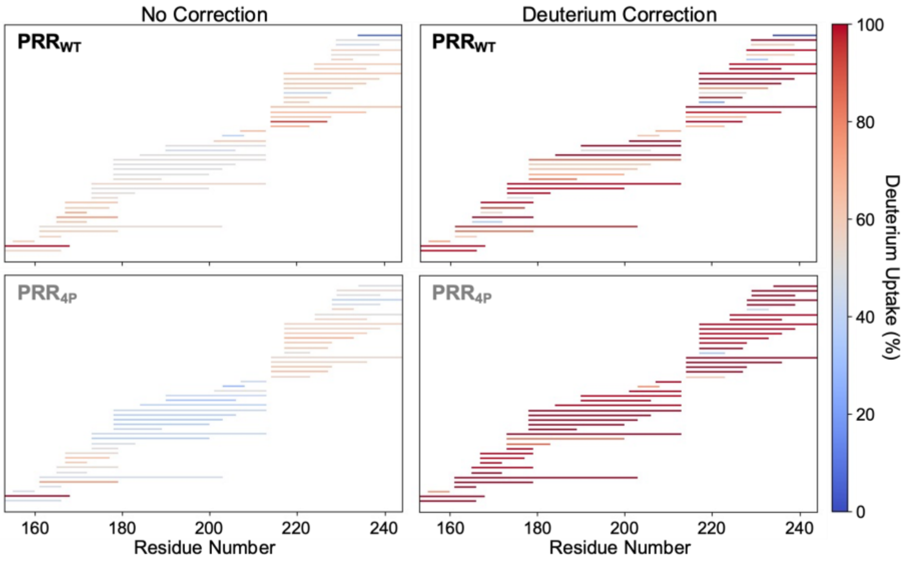


**Figure S9. Comparison of $PRR_{WT}$ and $PRR_{4P}$ exchange.** Deuterium uptake measurements for $PRR_{WT}$ (black, top) or $PRR_{4P}$ (gray, bottom) with and without deuterium back correction. Overall, the $PRR_{4P}$ peptides appear to have less exchange than the $PRR_{WT}$. Correction for deuterium back exchange was calculated as described in the *Materials & Methods*: Eqn 6, where the corrected deuterium uptake (in number of deuterons) is based on the change in the centroid mass between the non-deuterated, and maximally deuterated samples relative to the total possible amide backbone exchange sites for each peptide. When corrected, peptides from both PRR constructs show much higher percentages of exchange and are more comparable in extent of exchange. Data is presented as mean ± SD, for n ≥ 3 independent measurements.
